# APE1 Coordinates Its Disordered Region and Metal Cofactors to Drive Genome Surveillance

**DOI:** 10.64898/2025.12.28.696760

**Authors:** Donghun Lee, Subin Kim, Gyeongpil Jo, Juwon Kim, Jungmin Yoo, Jejoong Yoo, Ja Yil Lee, Gwangrog Lee

## Abstract

Efficient recognition of DNA lesions such as apurinic/apyrimidinic (AP) sites is essential for maintaining genome stability. Apurinic/apyrimidinic endonuclease 1 (APE1) is the primary eukaryotic AP endonuclease, yet how it identifies rare lesions among vast stretches of undamaged DNA remains incompletely understood. Using single-molecule imaging combined with molecular dynamics simulations, we reveal that APE1 employs a distinctive dual mechanism to search DNA. First, Mg²⁺ coordination at the active site neutralizes clustered negative charges, stabilizing electrostatic contacts during scanning. Second, its N-terminal intrinsically disordered region (IDR)—a feature conserved only in eukaryotic homologs but absent in prokaryotic ExoIII—not only forms DNA through transient IDR contacts but also engages continuous interactions via the unprecedented 177 arginine residue within the structured nuclease domain, thereby prolonging residence time and enabling long-range diffusion. Together, these two modules synergize to promote a sliding-based search strategy tailored to the complexity of eukaryotic genomes. Consistent with this model, IDR deletion restricts APE1 to 3D collisions, whereas IDR duplication enhances 1D scanning. Thus, APE1 exemplifies how structural disorder and metal-ion coordination integrate to enable long-range lesion recognition, highlighting an evolutionary innovation in eukaryotic DNA repair.

## Introduction

A fundamental question in protein–DNA interactions is how DNA-binding proteins efficiently locate specific targets within vast genomes ^1, 2^. Many DNA-binding proteins overcome this challenge by employing facilitated diffusion mechanisms that combine three-dimensional (3D) collisions with one-dimensional (1D) diffusion (sliding and/or hopping) and intersegmental transfer along the DNA backbone ^3–6^.

Intrinsically disordered regions (IDRs), which are highly enriched in DNA-binding proteins including transcription factors and DNA repair enzymes, have emerged as key regulatory elements in this dynamic search process ^7, 8^. Unlike globular domains, IDRs are highly flexible and contain clusters of charged residues ^9^, enabling them to establish transient and electrostatic interactions with DNA. This flexibility allows proteins to slide smoothly along the DNA helix, extending their residence time and enhancing the efficiency of target site sampling. IDRs can thus tune the balance between rapid diffusion and stable recognition, a property that has been supported by direct single-molecule observations in multiple DNA repair factors ^10–12^.

In this context, efficient and accurate detection of DNA lesions is essential for maintaining genomic stability ^13–16^. Apurinic/apyrimidinic (AP) sites, or abasic sites, are the most frequent types of DNA damage, with an estimated 10,000 to 20,000 lesions occurring per cell each day. Unless repaired, AP sites can obstruct transcription and replication and are associated with cancer and other diseases. These lesions are primarily resolved through the base excision repair (BER) pathway, where apurinic/apyrimidinic endonuclease 1 (APE1) acts as the central enzyme to cleave the DNA backbone at AP sites ^17–19^.

Although the catalytic mechanism of APE1 is well characterized, how it recognizes AP sites among the vast excess of undamaged DNA remains poorly understood ^20^. This question is particularly relevant in the context of growing evidence that DNA repair proteins employ 1D diffusion to efficiently scan for DNA damage. Studies on OGG1, Msh2–Msh6, Mlh1–Pms1, and XPC–RAD23B have revealed that repair proteins utilize facilitated diffusion to rapidly search for DNA lesions even on nucleosomal DNA while maintaining specificity for damaged sites ^3, 21^.

This work raises an important and unexplored question: does APE1 similarly use 1D diffusion, and if so, what role does its N-terminal IDR play in this process? APE1 comprises a structured C-terminal endonuclease domain and a conserved N-terminal IDR that is present in eukaryotic homologs but lacks in prokaryotes ^22, 23^. This IDR contains a nuclear localization signal and has been implicated in redox signaling, transcriptional regulation, and protein–protein interactions. The redox-active motif within this region is known to enhance the DNA-binding of redox-sensitive transcription factors such as AP-1, NF-κB, and HIF-1α by reducing cysteine residues in their DNA-binding domains ^24–26^. However, whether APE1’s IDR contributes directly to its lesion search activity has not yet been experimentally addressed. Furthermore, because APE1’s enzymatic activity depends on divalent metal ions, it remains unclear whether ions like Mg²⁺ play a dual role in catalysis and in modulating the dynamics of target search ^27, 28^.

To address these questions, we hypothesized that APE1 uses 1D diffusion to locate AP sites and that its disordered N-terminal region facilitates this process. Moreover, we investigated whether Mg²⁺ influences not only catalytic efficiency but also APE1’s DNA-scanning behavior. To test this hypothesis, we employed single-molecule imaging to directly visualize APE1’s motion along DNA ^29, 30^. By manipulating IDR length and assessing diffusion kinetics and AP-site targeting efficiency, we uncovered a previously unrecognized function of the IDR in promoting lesion search. This IDR function was further supported by complementary all-atom molecular dynamics (MD) simulations. Moreover, we found that Mg²⁺ enhances the efficiency of this process, suggesting a dual role in coordinating both enzymatic activity and target site localization. Together, these findings reveal a critical role for structural disorder in APE1’s mechanism of action and highlight how eukaryotic DNA repair enzymes are finely tuned for genome surveillance.

## Materials and Methods

### 1. DNA Preparation

#### 1-1. DNA cloning and site-directed mutagenesis

Ligation-independent cloning was employed to clone the full-length wild-type human *APE1* (hereafter referred to as APE1) gene into the pB3 vector, which contains a 6× His tag followed by a TEV cleavage site and the gene of interest. Correct insertion was confirmed via sequencing. All APE1 mutants were generated using site-directed mutagenesis. Primers specific for the desired mutations were designed to amplify the pB3-APE1 vectors via PCR. Post-PCR, DpnI (R0176S, New England Biolabs) digestion was performed to remove the original plasmid. The resulting PCR product was transformed into *E. coli* DH5α cells, and single colonies were picked for sequencing to confirm the presence of the intended mutations.

#### 1-2. Oligonucleotide Synthesis and Labeling

All oligonucleotide DNAs were synthesized by IDT, Bioneer, or Bionics. Oligonucleotides containing 5′-C6, internal amino-modifier C6 dT, or 5′-amino-modifier C6 groups were labeled with Cy3-NHS (PA13101, Cytiva) or Cy5-NHS (PA15101, Cytiva) at room temperature for 6 hours. Free dyes were removed via ethanol precipitation. Labeled ssDNA was annealed with complementary strands in 10 mM Tris-HCl (pH 8.0), 50 mM NaCl buffer by heating to 95°C for 3 minutes and cooling slowly to room temperature.

#### 1-3. λ-I3 DNA Preparation

λ-I3 (47,472 bp) is an engineered lambda phage DNA (λ-DNA) that contains seven nickase (Nt.BspQI) sites between 33,514 bp and 33,630 bp. λ-I3 was prepared by following the previous protocol ^60, 61^. The λ-DNA was amplified using MaxPlax λ extracts (MP5105, Epicentre). The phage particles containing λ-I3 were then infected into *E. coli* LE302MP strain. The infected *E. coli* cells were grown up at 37°C for 20 min and then spread on an agar plate with 0.7% of top agar. After overnight incubation at 37°C, a single plaque was taken and added to 800 µL of LE392MP cells supplemented with 5 mM MgCl_2_ and 5 mM CaCl_2_. The mixture with the single plaque was incubated in 200 mL of NZCYM broth at 37°C with 125 rpm. OD_600_ rose up over 1.6 and suddenly dropped down close to ∼0.3 because of bacteriophage burst. Immediately, 5 mL of chloroform was added to the cells followed by further incubation at 37°C for 15 min. NaCl powder was dissolved to the cells up to 1 M on ice for 10 min. The cells were pelleted at 12,000 g for 10 min, and the supernatant was collected. PEG20000 (81300, Merck) was added to the supernatant up to 10% (w/v) and incubated at 4°C at least overnight. The phage particles were spun down at 12,000 g for 15 min. The phage pellets were resuspended in SM buffer (10 mM Tris-HCl [pH 8.3], 100 mM NaCl, and 10 mM MgCl_2_). Then RNase A (R4875-500, Merck) and DNase I (D5319-2, Merck) were added to degrade genomic DNA and RNA, respectively. Proteinase K (P2308, Merck) was treated and incubated at 37°C for 45 min to remove RNase A and DNase I. Proteinase K was subsequently heat-inactivated at 65°C for 10 min with 1.25% of SDS and 100 mM EDTA. λ-I3 was finally purified by isopropanol precipitation.

For DNA curtain experiments, 10 nM of lambda DNA (N3011S, NEB) or λ-I3 was mixed with 1 μM of lambda R-biotin oligomer and 1 μM of lambda L oligomer in T4 DNA ligase buffer (M001S, Enzynomics). The mixture was heated at 65℃ for 10 min and then cooled down to 23℃. Then 40 units/μL of T4 DNA ligase was added and incubated at 23℃ overnight for sealing the nicks.

#### 1-4. AP Site Incorporation into λ-I3

Insertion of AP sequences into λ-I3 was conducted by following the previous protocols (Figure S7A) ^60, 61^. 6 units/mL of Nt.BspQI (R0644S, NEB) was treated to 10 nM of λ-I3 in 1× NEBuffer r3.1 (B6003S, New England Biolabs) with 1 mM ATP and then incubated at 50℃ for 1 hr. To inactivate nickases, proteinase K was added to the reactant, which was further incubated at 50℃ for 1 hr. 1 μM of lambda R-biotin oligomer, 1 μM of lambda L oligomer, and 5 μM of AP site oligomer were incubated with λ-DNA at 80℃ for 20 min and slowly cooled down to room temperature. During this process, a gap between 33,514 bp and 33,630 bp was hybridized with the excessive AP oligomer. All nicks were sealed by T4 DNA ligase (M001S, Enzynomics) at 23°C overnight. Unligated DNA oligomers in the mixture were removed by MicroSpin S-400 HR Columns (27514001, Cytiva). Because the AP-inserted region of λ-I3 contained NcoI cleavage sites, the AP insertion was confirmed by NcoI cleavage (Figure S7B). 0.5 nM AP-containing λ-I3 was treated with 1 units/μL NcoI (R0193S, NEB) at 37°C for 1 hr. The digested fragments were analyzed 1% agarose gel electrophoresis (Figure S7C).

### 2. Protein Expression, Purification, and Labeling

#### 2-1. Expression and Purification of APE1 and Mutants

Wild-type APE1 and its variants were purified following previously established protocols. All APE1 variants were expressed in BL21(DE3)-star *E. coli* cells (C601003, Thermo Fisher Scientific). Cells were cultured in LB medium containing 100 µg/ml ampicillin at 37°C until the OD600 reached 0.6. Protein expression was induced by adding 1 mM IPTG, followed by incubation at 16°C for 18 hours. Cells were harvested by centrifugation at 5,000 g for 10 min at 4°C and resuspended in lysis buffer (20 mM Tris-HCl [pH 7.5], 250 mM NaCl). Cell lysis was performed using sonication (10 sec pulse on, 20 sec pulse off, total 9 min). The lysate was clarified by centrifugation at 35,000 g for 30 min at 4°C and filtered through a 0.45 µm syringe-driven filter (Millipore).

APE1 was purified via nickel affinity chromatography using a HisTrap HP column (17524701, Cytiva) on an FPLC system. Protein was eluted with a gradient of buffer B (20 mM Tris-HCl [pH 7.5], 250 mM NaCl, 500 mM imidazole). Fractions containing APE1 were pooled, dialyzed against 20 mM Tris-HCl [pH 7.5], 250 mM NaCl, and 10 mM 2-mercaptoethanol, and supplemented with 20% glycerol before being aliquoted and snap-frozen in liquid nitrogen for storage at −80°C.

For labeling mutants, proteins were dialyzed in the same buffer and subjected to TEV protease cleavage to remove the His tag. The cleavage reaction mixture was passed through a second nickel column (17524701, Cytiva) to remove the His-tagged TEV protease and His-tag fragments. The flow-through containing APE1 was concentrated using Amicon Ultra-15 centrifugal filters (UFC901024, Merck Millipore), aliquoted with 20% glycerol, snap-frozen, and stored at −80°C.

Wild-type *APE1* gene and D210N mutant were cloned into pB3 plasmid, and 6x His and TEV protease site at the amino-terminus and 3× HA (hemagglutinin) tag at the carboxyl-terminus were inserted. The purification of APE1^D210N^ followed the same protocol as wild-type APE1. The protein was expressed in *E. coli* strain BL21(DE3). The cells were grown in 1 L LB broth supplemented with 1 mg/mL ampicillin at 37℃until OD_600_ was about 0.6. 0.5 mM IPTG (Isopropyl β-D-1thiogalactopyranoside) was added to induce protein expression, and the cells were further grown at 16℃ overnight. The cells were harvested by centrifugation at 8,000 g and resuspended in 10 mL of APE1_Resuspension buffer (20 mM Tris-HCl [7.5], 250 mM NaCl, and 1 mM PMSF). After lysed by sonication, the cell lysates were ultracentrifuged at 100,000 g for 1 hr. Then the clarified lysates were loaded onto HisTrap HP 1 mL column (Cytiva, 29051021) that was equilibrated with 5 mL of APE1_Washing buffer (20 mM Tris-HCl [7.5] and 1 M NaCl). In all purification steps, the flow rate for HisTrap HP 1 mL column was 0.5 mL/min. The column was washed with 5 mL of APE1_Washing buffer. The protein was eluted by imidazole gradient from 0 mM to 500 mM in APE1_Resuspension buffer. The protein was usually eluted at approximately 200 mM imidazole. Fractions containing APE1 were pooled, and his-TEV protease (1:50 w/w, New England Biolabs, P8112S) was added to remove 6x His tag. The sample was then dialyzed against APE1_Dialysis buffer (Tris-HCl [7.5], 250 mM NaCl, and 5 mM 2-mercapto-ethanol) overnight. The dialyzed sample was loaded onto HisTrap HP 1 mL column that was equilibrated by APE1_Washing buffer and flow-through were collected to remove 6x His tag and his-TEV protease. Finally, purified APE1-3xHA was stored at −80℃ ^39, 62^.

#### 2-2. Protein labeling

For smFRET experiments, site-specific N-terminal labeling of APE1 was achieved using a calcium-independent sortase. TEV cleavage exposed a 6x glycine sequence at the N-terminus of APE1. The labeling reaction was carried out at room temperature for 30 min in a buffer containing 50 mM Tris-HCl [pH 7.5] and 150 mM NaCl, using 10 µM APE1, 20 µM sortase, and 100 µM Cy3-LPETGG probe (purchased from ANYGEN). Unreacted Cy3-LPETGG probe was removed using Zeba desalting columns (89882, Thermo Scientific™), followed by passage through a nickel resin to remove His-tagged sortase. Purified Cy3-APE1 was analyzed by 12% SDS-PAGE and visualized using a ChemiDoc XRS+ system and Coomassie blue staining.

For DNA curtain experiments, APE1 was labeled with Cy5. The ybbR-tagged C-terminus of APE1, Sfp proteins, and CoA-conjugate Cy5 were mixed at a molar ratio of 1:2:10 in 1x Sfp reaction buffer (50 mM Tris-HCl [pH 7.5] and 10 mM MgCl_2_) and incubated for 30 min at 23℃. Then, the reactant was loaded onto 5 mL of HisPur^TM^ Ni-NTA resin (88221, Thermo Fisher Scientific) which was equilibrated with 5 mL of Cy5-APE1 Washing buffer (20 mM Tris-HCl [pH 7.5] and 250 mM NaCl). The column was washed with Cy5-APE1 Washing buffer. The protein was eluted with Cy5-APE1 Elution buffer (20 mM Tris-HCl [pH 7.5], 250 mM NaCl, and 100 mM imidazole). Fractions containing Cy5-labeled APE1 were pooled and then dialyzed against APE1_Dialysis buffer. The labeling efficiency was estimated from the ratio of absorption of Cy5 and APE1 using Nanodrop (Thermo Fisher Scientific).

### 3. Biochemical Assay

#### 3-1. Enzyme activity assay

The activity of purified APE1 was tested from the cleavage of DNA containing an AP site. 10 nM of Cy5-labeled 25-nt AP DNA was mixed with different concentrations of APE1 (0 nM ∼ 300 nM) in 1× APE1 cleavage buffer (20 mM Tris-HCl [pH 7.5], 100 mM NaCl, 0.1 mM MgCl_2_, and 1 mM DTT). The mixtures were incubated at 23℃ for 5 min. After reaction, the same volume of formamide denaturing dye (95% formamide and 40 mM EDTA) was added to the mixtures and further incubated at 95°C for 5 min. The reactants were analyzed by 15% urea denaturing gel electrophoresis ^55, 63^.

#### 3-2. Electrophoretic mobility shift assay (EMSA) for AP DNA

To test the binding affinity^64^ of APE1 to AP DNA, 10 nM Cy5-labeled 25-nt AP DNA (or 25-nt homoduplex DNA) was mixed with different concentrations of APE1 (10 nM ∼ 500 nM) in 1× APE1 binding buffer (20 mM Tris-HCl [pH 7.5], 100 mM NaCl, 1 mM EDTA and 1 mM DTT). The mixtures were incubated at 23℃ for 5 min. The reactants were analyzed by 6% non-denaturing polyacrylamide gel electrophoresis ^35, 55, 63^.

#### 3-3. EMSA for undamaged DNA

EMSA was conducted by incubating 2 nM DNA with 0∼1000 nM APE1 in 20 mM Tris-HCl [pH 7.5], 0.5 mM EDTA, 10 mM NaCl, 1 mM DTT, 10% glycerol. Complexes were separated on 15% native PAGE and imaged using ChemiDoc (Bio-Rad).

### 4. Single-Molecule FRET (smFRET) Analysis

#### 4-1. Microscope Setup and Sample Preparation

smFRET was performed on a prism-based TIRF microscope (Olympus IX71). Quartz slides and coverslips were PEG-passivated. Biotinylated DNA was immobilized via biotin–neutravidin linkage. Excitation was achieved with a 532-nm laser; emissions were split by a 630-nm dichroic and captured using an EMCCD (iXon Ultra 897, Andor). FRET was calculated using:

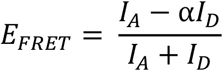

where *I*_*D*_ and *I*_*A*_ are background-corrected donor and acceptor intensities, respectively, and *α* accounts for donor leakage. A 532-nm laser provided TIRF illumination, exciting Cy3, and the emission was collected using a 60× water immersion objective. Data acquisition was performed at 100 msec frame intervals and analyzed using IDL, MATLAB, and Origin.

#### 4-2. Binding and Dissociation Assay

Biotin-DNA (50 pM) was immobilized and equilibrated in imaging buffer (50 mM Tris-HCl [pH 7.5], 100 mM NaCl, BSA, DTT, oxygen scavengers). 0.5 nM Cy3-APE1 was injected and monitored at 100–500 msec intervals. Individual binding trajectories were analyzed to count translocation events per binding.

3-3. EMSA for Undamaged DNA

EMSA was conducted by incubating 2 nM DNA with 100 nM APE1 in 20 mM Tris-HCl [pH 7.5], 0.5 mM EDTA, 10 mM NaCl, 1 mM DTT, 10% glycerol. Complexes were separated on 15% native PAGE and imaged using ChemiDoc (Bio-Rad).

### 5. DNA Curtain Assay

All DNA curtain experiments were conducted at 23°C following by previous protocols ^65, 66^.

#### 5-1. Preparation of double-tethered DNA curtain assay

A fused-silica slide with nano-trench patterns was assembled into a flowcell containing a microchamber. 0.5% biotinylated-DOPE (1,2-dioleoyl-sn-glycero-3-phosphoethanolamine-N-(cap biotinyl) (sodium salt), 870273, Avanti Polar Lipids), 8% mPEG 2000-DOPE (1,2-dioleoyl-sn-glycero-3-phosphoethanolamine-N-[methoxy(polyethylene glycol)-2000], 880130, Avanti Polar Lipids) and DOPC (1,2-dioleoyl-sn-glycero-3-phophocholine, 850375, Avanti Polar Lipids) were mixed to compose liposomes, which were injected into the flowcell to form lipid bilayer on the slide surface. 0.1 mg/mL of anti-digoxigenin (11214667001, Roche) was absorbed on the pentagonal nano-structure by 40-min incubation. The slide surface was further passivated with 0.4% BSA (bovine serum albumin, A7030, Sigma). Then 25 μg/mL of streptavidin was injected. After free streptavidin was removed, 50 pM biotinylated λ-DNA were anchored on lipid bilayer via biotin-streptavidin linkage. Under 4 mL of continuous flow at 0.3 mL/min flow rate, digoxigenin-modified end of λ-DNA was tethered to the pentagonal nano-barrier, and hence DNA remained stretched even in the absence of buffer flow.

#### 5-2. DNA curtain experiments for APE1

All DNA curtain experiments for APE1 were conducted in APE1 buffer (20 mM Tris-HCl [pH 8.0], 0.05 mg/mL BSA, and 1 mM DTT) supplemented with different NaCl and MgCl_2_ concentrations at 23℃. For fluorescence imaging, APE1 was labeled with HA-antibody-conjugated quantum dot with 705 nm emission (Qdot-705) (S10454, Thermo Fisher Scientific). APE1 and HA-antibody-conjugated Qdot-705 were mixed at 1:6 molar ratio. The mixture was incubated on ice for 15 min. 0.3 to 2 nM APE1 was injected into the flowcell. When a maximum amount of APE1 arrived at DNA curtains, the flow was turned off. Subsequently, Qdot fluorescence signal was imaged by NIS-Element software (Nikon) under the illumination of 488-nm laser (200 mW, OBIS, Coherent Laser) with 0.1 sec exposure time for 5 min.

#### 5-3. Data analyses for DNA curtain experiments

All images were converted into TIFF format and processed by ImageJ (NIH). For APE1 molecules showing DNA binding after the flow was stopped, the initial binding position of APE1 on lambda DNA was determined by fitting a two-dimensional Gaussian function to the Qdot fluorescence signal and extracting its center. The error was obtained from bootstrapping with 70% confidence interval.

#### 5-4. 1D diffusion analysis

To observe 1D diffusion of APE1 along DNA, APE1 was incubated in the flowcell for 5 min. After unbound APE1 was removed, the flow was turned off, and the movement of APE1 was imaged. The motion of APE1 was tracked by an ImageJ plug-in, MOSAICSuite particle tracker. 1D diffusion coefficient (*D*) was calculated from mean square displacement (*MSD*) as previously described ^10, 67^ The MSD equation was given as

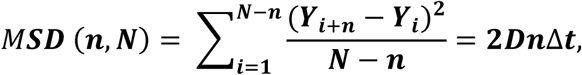

*N*: total number of frames.

*n*: measurement window ranging from 1 to N-1.

*Δt*: time interval between frames

*Yi*: APE1 position at *i*^th^ frame.

Error of *MSD* was calculated using standard deviation (SD) given as

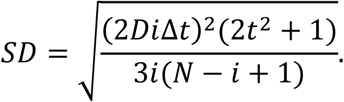

After linear fitting of MSD with first three data points, the diffusion coefficient (D) was obtained from the slope.

#### 5-5. Calculation of theoretical limit of diffusion coefficient for rotation around DNA helix

The diffusion limit of rotational motion around DNA helix was calculated based on modified Schurr’s model as previously described ^68^. The total friction was given like

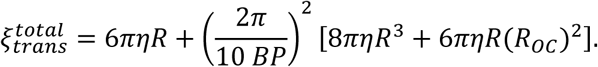

Here, *η* is the viscosity of water (0.9 mPa sec), *BP* is the length of 1 bp of duplex DNA (0.34 nm), *R* is the radius of protein, and *R_OC_* is the separation between DNA axis and the center of mass of protein. Then the theoretical limit of diffusion coefficient for the rotational motion is given from Einstein equation,

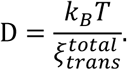

APE1 consists of intrinsically disordered domain at the amino-terminus and glycosylase domain. The radius of glycosylase domain is estimated from the crystal structure as 2.53 nm ^69–71^. The intrinsically disordered region contributes to the radius, which is estimated from Kuhn length of freely joint model as below,

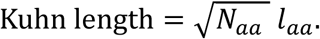

Here, *N_aa_* is the number of amino acids for the intrinsically disordered region, which is 64 for APE1. *l_aa_* is the length between amino acids, which is 0.432 nm. Therefore, the radius of intrinsically disordered region is estimated as 3.456 nm. The total radius of APE1 is given as 5.983 nm. On the other hand, APE was labeled with Qdot in the DNA curtain assay, and hence *R_OC_* is calculated as the sum of APE1 total radius and the hydrodynamic radius of conjugated antibody. The hydrodynamic radius of antibody-conjugated Qdot is ∼ 13 nm based on the information of maker (Thermo Fisher Scientific). So, *R_OC_* is approximately 20 nm. *k_BT_* is approximated as 4.1 pN nm at 23°C. Finally, the rotational limit of diffusion coefficient of APE1 is ∼ 0.004 μm^2^/sec, which is smaller than the diffusion coefficients of APE1 at all NaCl concentrations. For Cy5-labeled APE1, the radius of Cy5 is given as 0.82 nm. Then the rotational limit of diffusion coefficient is ∼0.076 μm^2^/sec. Most diffusion coefficients of Cy5-APE1 are higher than 0.076 μm^2^/sec. There results suggest that APE1 translationally slides along the DNA backbone without rotation around the helix.

#### 5-6. Lifetime and AP-residence time analyses

To analyze the lifetime of diffusing APE1 on single DNA, kymographs were drawn for individual DNA molecules. The time intervals between binding and dissociation on DNA were collected from kymographs and fitted by a single exponential decay function for the lifetime. For AP-residence time, APE1 molecules that bound to AP sites were chosen. The binding time intervals at AP sites were collected from kymographs and then fitted by a single exponential decay function for AP residence time. In some cases, because we did not observe the dissociation, the AP residence time was underestimated.

### 6. MD simulation

#### 6-1. General MD simulation protocol

All-atom molecular dynamics (MD) simulations were performed with GROMACS 2023.1 ^49^ using the AMBER ff99sb-ildn-phi force field ^45–47^ with the parmbsc0 modification for nucleic acids ^72^ and the CUFIX corrections for non-bonded interactions ^42, 48^ and the original TIP3P water model ^73^. Following energy minimization, systems were equilibrated in the NVT and NPT ensembles for 100,000 steps each with a 1 fsec integration timestep. Production runs employed a 2 fsec timestep under periodic boundary conditions applied in all three dimensions. Short-range nonbonded interactions were treated using the Verlet cutoff scheme, with a 1.2 nm cutoff for both van der Waals and Coulomb interactions. A switching function was applied to the van der Waals potential between 1.0 and 1.2 nm. Long-range electrostatics were computed using the Particle Mesh Ewald method with a 1.2 nm real-space cutoff ^74^. All bonds involving hydrogen atoms were constrained using the LINCS ^75^ or the SETTLE ^76^ algorithms. The system temperature was maintained at 310 K using the velocity-rescale thermostat ^77^, while pressure was controlled semi-isotropically at 1 bar using the C-rescale barostat ^78^.

#### 6-2. MD simulation setup

We obtained the atomistic coordinates of APE1 without IDR from the crystal structure (PDB: 1DEW) ^17^. The IDR (residues 1 to 40) in an extended form was reconstructed using AlphaFold3 ^43^. For both types of APE1, we generated four independent systems, and the protein’s initial position was randomized in each replicate. The system comprises one DNA strand and one APE1 molecule, placed in the hexagonal box (*a = b ≈ 10.1 nm, c ≈ 10.5 nm*, *α* = *β* = 90°, *γ* = 60°) at 100 mM NaCl concentration. The DNA featured a 32-base-pair repeating CG sequence, and we aligned the DNA axis to the z-axis and covalently bonded two ends of a DNA strand to approximate an effectively infinite DNA ^44^. The average simulation time for the four native APE1–DNA systems was about 40 μsec, whereas for the tail-truncated systems, the average simulation time was about 10 μsec.

#### 6-3. MD simulation analysis

Before analysis, we removed DNA’s translational displacement by fitting each frame’s DNA atoms to their initial coordinates using GROMACS’s trjconv tool, recording atom positions every 1 nsec. We then applied the same tool to correct any discontinuities when APE1 crossed a periodic boundary and wrapped to the opposite side of the box, thereby preserving a continuous trajectory. Next, for each frame, we measured the distance between APE1’s center of mass and the DNA axis, assigning the index of the closest DNA. Hopping events were identified at points where peaks in the APE1-DNA distance trace coincided with step changes in the DNA-index plot. We then used the trjconv tool to split the trajectory at each hopping event, adjusting frames to ensure that APE1 remained bound to the nearest DNA and positioned within the central unit cell. To eliminate residual rotational displacements, we superposed the DNA atoms of every frame onto those of the initial structure using the MDtraj Python package ^79^. With the cleaned trajectories in hand, we converted APE1’s center-of-mass coordinates into cylindrical form (*z*, *r*, *θ*) and computed both the translational mean squared displacement of *z*(*t*) and the rotational mean squared displacement of *θ*(*t*) as functions of the lag time *Δt*. Finally, we quantified APE1-DNA contacts on a per-residue basis. Residues with the terminal atom (Nz for Lys; Cz for Arg; Cb for Ala) located within 1.25 nm from the DNA axis were classified as “in contact,” otherwise “no contact,” producing a binary contact matrix (1 = contact, 0 = no contact). We then averaged over the time axis to calculate the fraction of contacts for each residue. We also summarized contacts at the domain level by marking the structured domain as “in contact” if at least one binding-site residue was in contact, and the IDR (tail) as “in contact” if any residue from residue number 1 through 11 was in contact.

## Results

### One-dimensional diffusion of APE1

To investigate the interaction of APE1 with DNA, we employed a single-molecule Förster resonance energy transfer (smFRET) assay (Figures 1A and S3). The donor fluorophore (Cy3) was site-specifically conjugated to the N-terminus of APE1 using a sortase-mediated labeling method, and Cy3-labeled APE1 was directly visualized by excitation during its interaction with target DNA. We first examined APE1 binding using a DNA duplex containing a single AP site. The Cy5 as an acceptor was positioned 10 bp downstream of the AP site to detect FRET upon binding of Cy3-labeled APE1 (Figure S3). Upon binding of APE1 to the AP site, the proximity between the donor and acceptor dyes produced a stable FRET signal (∼0.38), indicative of a bound state (Figure S3B and S3C). Dissociation of APE1 from the DNA resulted in an abrupt loss of the FRET signal, marking a transition to the unbound state (Figure S3B). Notably, no binding events were observed on undamaged DNA, confirming the specificity of APE1 for AP sites.

**Figure 1.**
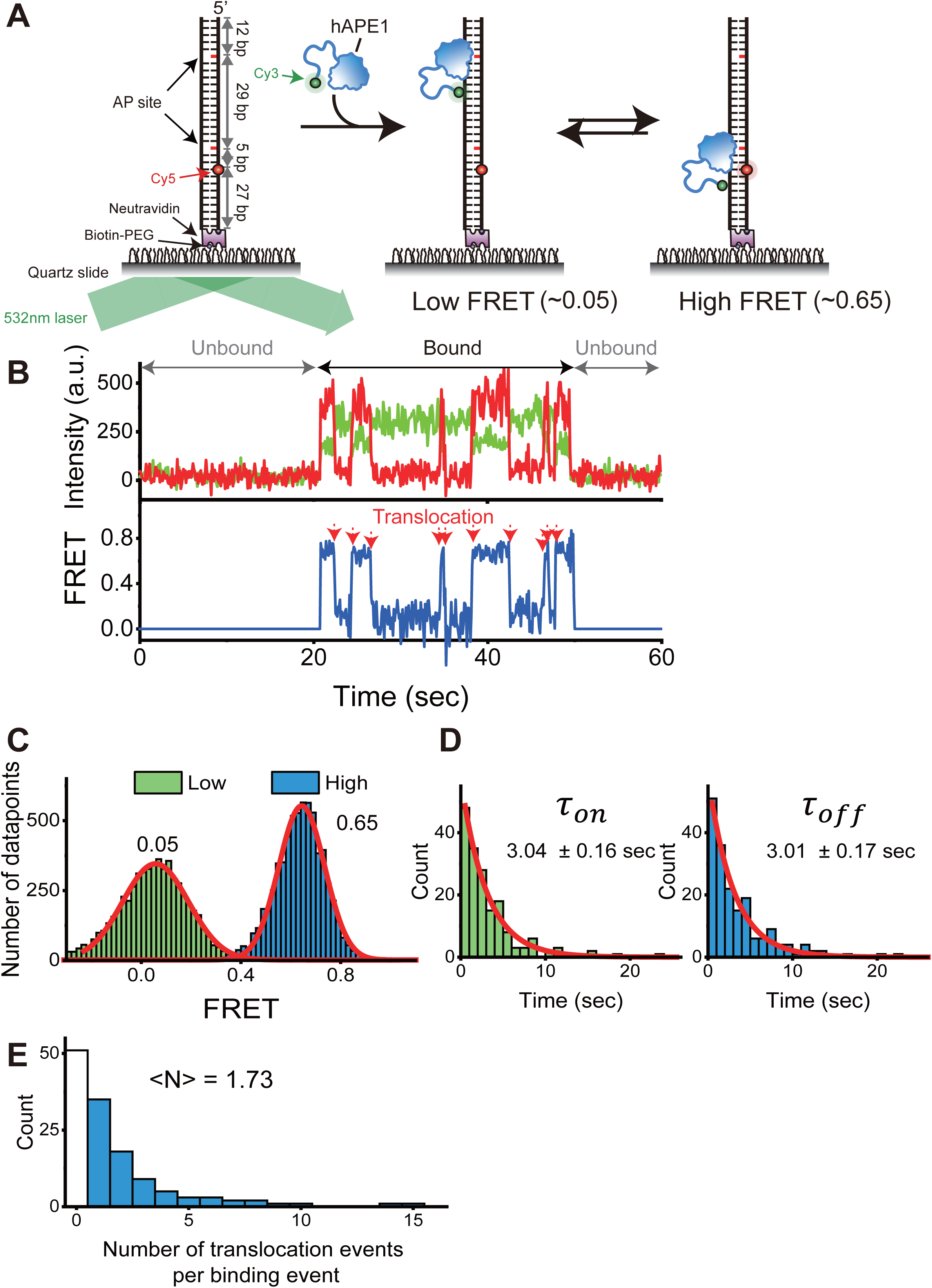
smFRET assay to monitor APE1 diffusion on DNA. (A) Schematic representation of the smFRET assay showing states before binding (left), after binding (middle), and during translocation between two AP sites (right) on a Cy5-labeled 2AP DNA substrate with Cy3-labeled APE1. (B) Representative smFRET trajectory showing APE1 binding and translocation events. Red arrows indicate rapid FRET transitions from low to high or high to low FRET states, occurring within a single frame (<100 ms). These rapid transitions represent APE1 translocation between two AP sites on the same DNA molecule. (C) FRET histograms of APE1 binding, categorized into low-FRET (green) and high-FRET (blue) states. (D) Histograms of binding durations for low-FRET (left, n = 223) and high-FRET (right, n = 148) states. (E) Distribution of translocation events per binding event (bin size = 1). APE1 undergoes an average of 1.73 shuttling events before dissociation. The white bar indicates binding events followed by immediate dissociation.

To explore whether APE1 exhibits local 1D diffusion along DNA, we next employed a substrate containing two AP sites separated by 29 bp, with the Cy5 acceptor dye positioned 5 bp downstream of the lower (bottom) AP site (Figure 1A). This dual-AP-site construct enabled us to detect APE1 binding and movement between distinct lesion sites based on FRET signal changes. However, it should be noted that we could only detect the bound state, not progressive transitions, due to the limited temporal resolution of our imaging system (100 msec), which is insufficient to capture the μsec-scale diffusion of APE1 along DNA.

This dual-AP-site construct revealed dynamic binding behavior consistent with local diffusion. When APE1 bound to the upper AP site, the donor and acceptor dyes remained spatially distant, yielding a low FRET state (∼0.05). By contrast, binding to the lower AP site brought the dyes into proximity, producing a high FRET signal (∼0.65) (Figures 1B and 1C). In this transition, the low FRET state did not indicate dissociation of APE1 since the Cy3 fluorescence remained stable throughout, confirming the continued presence of the Cy3-labeled APE1. The FRET histogram displayed two distinct states corresponding to these binding positions, with nearly identical dwell times (∼3.0 sec) for each site (Figure 1D), supporting the conclusion that APE1 binds specifically to AP sites. In many binding events, we observed APE1 translocating between the two AP sites prior to dissociation, as evidenced by transitions between high and low FRET states without signal loss. On average, APE1 performed approximately 1.73 translocation events per binding interaction, as determined by calculating the mean number of transitions per binding trace (Figure 1E). Together, these results indicate that APE1 is capable of local 1D diffusion along DNA, dynamically scanning between closely spaced AP sites before dissociating. The smFRET platform thus provides a powerful and sensitive method for elucidating the dynamic binding and search mechanisms of APE1 at single-molecule resolution.

### Effect of Mg²⁺ ions on 1D diffusion of APE1 along DNA

APE1 requires Mg²⁺ ions as essential cofactors for its enzymatic activity. In the crystal structure of the APE1–AP-nicked DNA complex (PDB: 5DFF), a single Mg²⁺ ion is coordinated by acidic residues—D70, E96, and D308—with E96 directly interacting with the ion (Figure 2A) ^28^. This coordination is crucial for stabilizing the enzyme’s active site and facilitating its phosphodiester bond cleavage during base excision repair. While the catalytic role of Mg²⁺ has been well established, its potential involvement in APE1’s target search dynamics remains less explored. Given the importance of rapid and efficient lesion recognition in maintaining genome integrity, we sought to determine whether Mg²⁺ also contributes to APE1’s ability to perform 1D diffusion along the negatively charged phosphate backbone of DNA.

**Figure 2.**
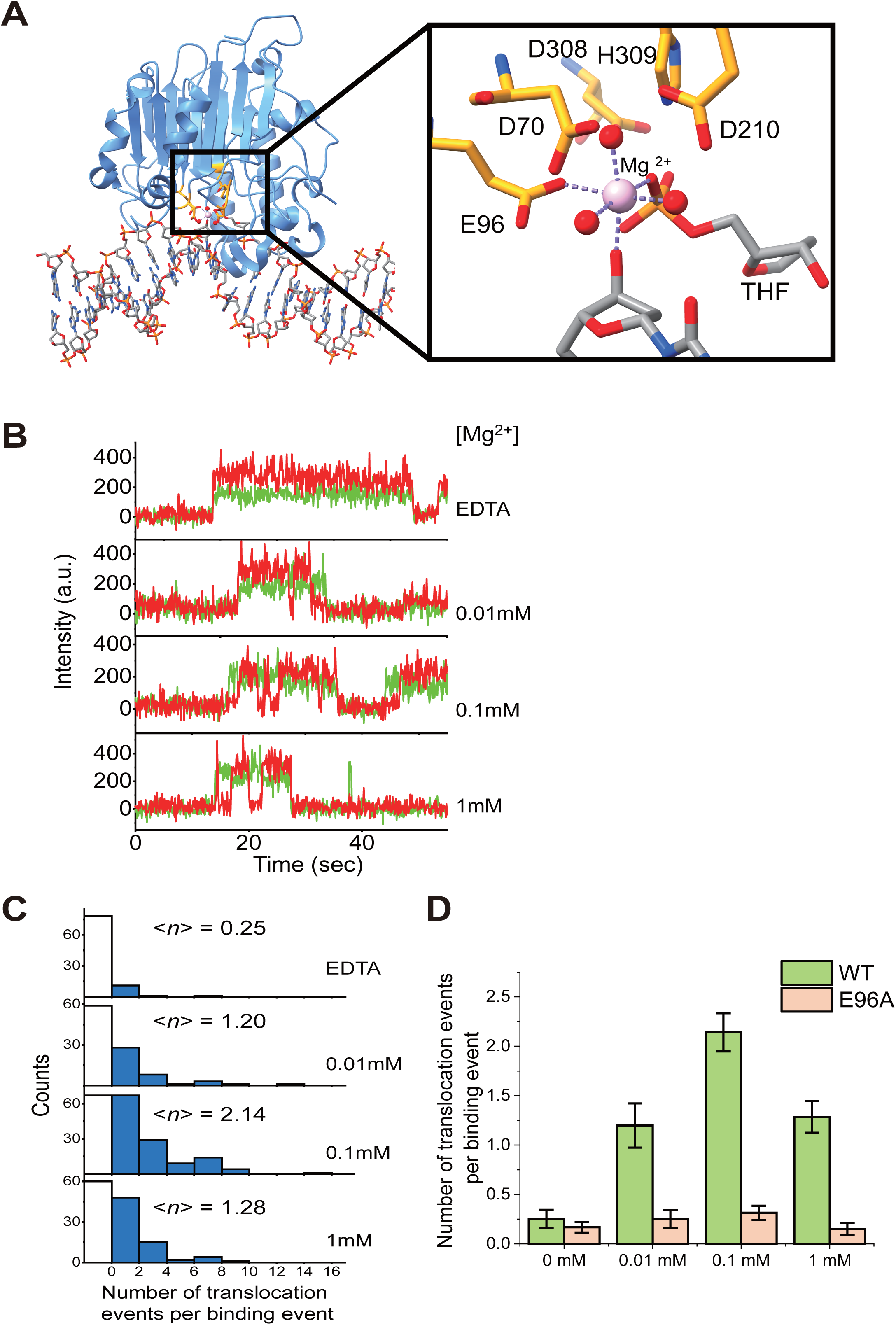
Effect of Mg²⁺ on APE1 diffusion. (A) Structural model of the APE1–AP site DNA complex showing Mg²⁺ coordination by residue E96. (B) Representative smFRET trajectories at varying Mg²⁺ concentrations. (C) Distribution of translocation events per binding at different Mg²⁺ levels (n = 91, 101, 191, 130 for EDTA, 0.01, 0.1, 1 mM, respectively). (D) Comparison of translocation frequencies between wild-type and E96A mutant APE1 under varying Mg²⁺ concentrations. Error bars indicate the standard error of the mean (s.e.m.)

To assess the effect of Mg²⁺ on APE1’s diffusion, we quantified the frequency of translocation events between two AP sites under varying Mg²⁺ concentrations. In the presence of EDTA without Mg²⁺, APE1 showed minimal translocation, averaging only 0.25 translocation events per binding (Figure 2B and 2C). In contrast, the addition of Mg²⁺ significantly enhanced translocation activity, peaking at 0.1 mM—a concentration close to physiological levels (Figures 2B and 2C) ^31–33^. These results suggest that Mg²⁺ facilitates not only catalytic cleavage but also the dynamic scanning behavior essential for efficient target search. To further elucidate the mechanism by which Mg²⁺ promotes diffusion, we examined the role of E96, a residue directly coordinating the Mg²⁺ ion. An E96A substitution mutant was generated to disrupt Mg²⁺ coordination without altering the overall structure of APE1. The E96A mutant exhibited AP site binding times comparable to the wild-type (Figure S4A–S4B), confirming that Mg²⁺ coordination via E96 does not affect static binding affinity. However, its translocation activity remained low (∼0.25 events per binding) regardless of Mg²⁺ concentration (Figure 2D), indicating a specific requirement for E96-mediated Mg²⁺ coordination in facilitating 1D diffusion.

These findings collectively reveal that Mg²⁺ enhances APE1’s local diffusion along DNA through a mechanism dependent on direct interaction with E96. Thus, Mg²⁺ plays a dual role: stabilizing the active site for catalysis and promoting dynamic movement along DNA to accelerate lesion recognition. This dual functionality highlights that Mg²⁺ is a critical cofactor to contribute to both efficiency and precision of APE1 in DNA repair.

### Effect of the IDR on the diffusion of APE1 along DNA

APE1 contains an intrinsically disordered region (IDR) at its N-terminus, which includes a nuclear localization signal (NLS) and has been implicated in multiple cellular functions such as DNA repair, redox regulation, and protein–protein interactions ^24–26^. In contrast to prokaryotic *E. coli* ExoIII, which lacks such disordered extensions, eukaryotic AP endonucleases universally possess this N-terminal IDR. This evolutionary divergence suggests that the presence of the IDR may provide a functional advantage, potentially compensating for the increased complexity and size of eukaryotic genomes. While smaller genomes may permit efficient target search via short 1D diffusion along DNA and/or 3D diffusion alone, the vast genomic landscape in eukaryotes likely demands more refined search mechanisms.

Given the growing evidence that facilitated diffusion crucial for efficient lesion recognition, we hypothesized that APE1’s IDR enhances its target search efficiency by promoting local diffusion along DNA. To evaluate this possibility, we engineered two APE1 mutants: one lacking the IDR entirely (0× IDR) and another with a tandem duplication of the IDR (2× IDR) (Figure 3A). We then quantified their translocation behaviors using single-molecule assays. Strikingly, deletion of the IDR (0× IDR) resulted in a 7.6-fold reduction in translocation events relative to wild-type APE1 (1× IDR), whereas the 2× IDR mutant exhibited a 1.5-fold increase in translocation frequency (Figures 3B and 3C). Despite these differences in dynamic movement, the binding dwell times across all constructs remained similar (Figure 3D), indicating that the IDR does not significantly alter binding affinity for AP sites, but rather modulates the ability to scan DNA through short-range diffusion.

**Figure 3.**
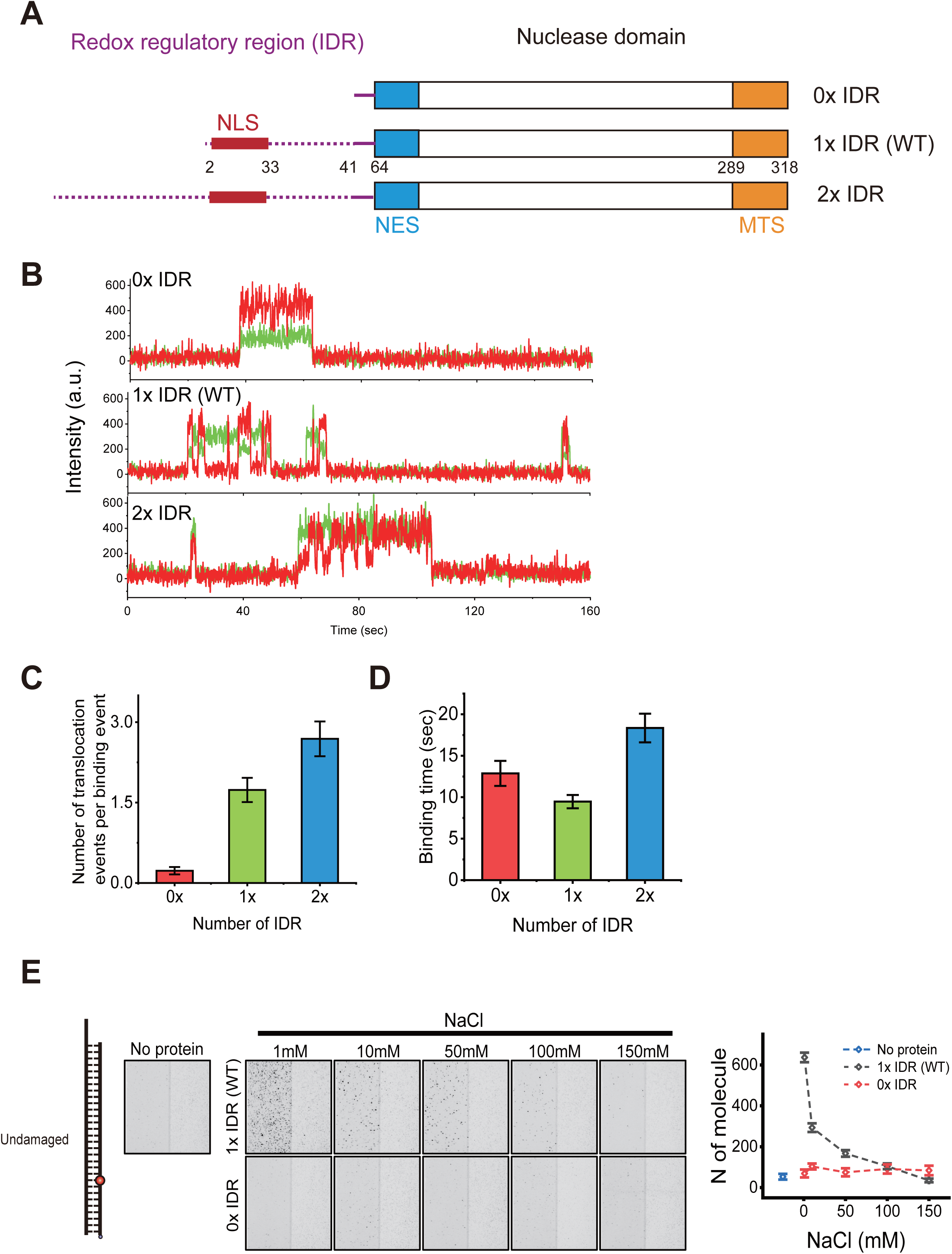
Role of the IDR in APE1 diffusion. (A) Domain architecture of APE1 constructs: wild-type (1× IDR), IDR-deleted (0× IDR), and IDR-duplicated (2× IDR). The redox regulatory region (IDR) is shown in purple. The structured nuclease domain is indicated in blue, white, and orange. NLS (nuclear localization signal) is shown in red; NES (nuclear export signal) in blue; and MTS (mitochondrial targeting sequence) in orange. (B) Representative smFRET trajectories showing the impact of IDR length on translocation behavior. (C) Quantification of translocation events per binding for each construct. Error bars represent s.e.m. (D) Binding lifetimes for wild-type and mutant APE1. Error bars indicate s.e.m. (E) Binding events of wild-type and 0× IDR APE1 to undamaged DNA at different NaCl concentrations (1–150 mM), quantified by the number of bound molecules with error bars representing s.e.m.

To further investigate whether the IDR contributes to nonspecific DNA interactions, we assessed APE1 binding to undamaged DNA under increasing NaCl concentrations. Wild-type APE1 exhibited measurable binding that decreased with higher salt concentrations, consistent with electrostatically driven, transient DNA interactions (Figure 3E). In contrast, the 0× IDR mutant failed to bind undamaged DNA under any tested condition. These observations were further supported by EMSA showing that wild-type APE1 formed DNA–protein complexes at higher protein concentrations, whereas the 0× IDR mutant did not (Figure S5).

Together, these results demonstrate that the N-terminal IDR enables APE1 to establish transient, nonspecific contacts with DNA, thereby preventing premature dissociation and facilitating local diffusion. This enhancement of 1D diffusion likely plays a critical role in efficient lesion search, particularly within the complex chromatin environment of eukaryotic cells.

### Long-range 1D diffusion of APE1 along extended DNA

Our smFRET data suggest 1D diffusion of APE1 for AP site search. However, human genome consists of approximately three billions of base pairs. To investigate how APE1 searches for AP sites in such long DNA, we adopted the single-molecule DNA curtain assay^34^ to directly visualize the motion of APE1 on DNA (Figure 4A). Wild-type and D210N mutant of APE1 with 6x His tag and TEV tag at the amino (N)-terminus and 3x HA tag at the carboxyl (C)-terminus were purified (Figure S1). The activity of purified APE1 was tested by nick generation at an AP site, which was observed by cleavage of DNA oligomer using denaturing gel electrophoresis (Figure S2A). Wild-type APE1 rapidly made a nick at the AP site, whereas D210N mutant did not. We next tested DNA binding affinity of both wild-type and D210N mutant in the absence of Mg^2+^ using EMSA and smFRET (Figure S2B-SE). D210N mutant showed higher binding affinity to an AP site than wild-type APE1, which is consistent with the previous report ^35^. However, either of them barely bound to undamaged DNA (Figure S5). For fluorescence imaging in DNA curtains, APE1 was labeled with HA-antibody-conjugated Qdot with 705 nm emission (S10454, Thermo Fisher Scientific). To avoid multiple APE1 from binding to single Qdot, APE and Qdot were mixed at the 1: 6 molar ratio. We first examined the behavior of APE1 on undamaged DNA (Figure 4A). APE1 initially bound to DNA without any sequence preference, indicating that the initial binding of APE1 to DNA is sequence-independent (Figure 4B). Then APE1 moved on DNA without any special orientation, showing that APE1 searches for AP sites through 1D diffusion along DNA (Figure 4C). APE1 mostly exhibited diffusive motion (86%) while a small portion of APE1 (14%) was immobile on DNA nonspecifically (Figure 4D). We next examined how long APE1 diffuses along DNA depending on ionic strength. The diffusion lifetime was determined as the time interval between the initial binding of the protein to DNA and its eventual dissociation, during which the protein displays 1D diffusion behavior along the DNA. The diffusion lifetime monotonically decreased as Na^+^ concentration increased, suggesting that the binding between APE1 and DNA is mediated by electrostatic interaction (Figure 4E and Figure S8A-S8D). These results are consistent with smFRET data (Figure 3E). On the other hand, for Mg^2+^, APE1 exhibited the longest lifetime at 0.1∼0.3 mM Mg^2+^, which is comparable to physiological Mg^2+^ concentration ^31–33^. Consistently with smFRET data, DNA curtain data show that Mg^2+^ plays a critical role in the diffusion of APE1 along DNA by maintaining APE1’s contacts with DNA under the physiological condition (Figure 4F and figure S8E-S8J). The diffusive motion of individual APE1 molecules was analyzed using single particle tracking (Figure 4C and Figure S6). The distributions of relative displacements were well fitted by single Gaussian functions, demonstrating that APE1 diffusion follows Brownian motion (Figure S6A). We estimated diffusion coefficients (D) by linear fitting of the first three data points of mean square displacement (MSD) derived from each time trajectory (Figure S6B). The diffusion coefficients were insensitive to NaCl and MgCl_2_ concentrations (Figure 4G and 4H). This independence of ion strength suggests that APE1 slides along DNA rather than hops during 1D diffusion ^10, 36^. Moreover, the theoretical limit of diffusion coefficient for the rotation around DNA helix of Qdot-labeled APE1 was estimated as 0.004 μm^2^/sec, which is lower than most diffusion coefficients of APE1, suggesting that APE1 slides along DNA without helical rotation (Figure 4G and 4H) ^36^. As the hydrodynamic diameter of Qdot is approximately 26 nm, we examined if Qdot influences the diffusion of APE1 using Cy5-labeled APE1. Cy5-labeled APE1 also showed the diffusive motion, which was analyzed in the same manner as Qdot-labeled APE1 (Figure S6C-S6F). The diffusion coefficients of Cy5-labeled APE1 were similar to those of Qdot-labeled APE1, suggesting that Qdot does not affect the diffusive motion (Figure S6G). Even though the theoretical limit of diffusion for helical rotation was increased (∼ 0.076 μm^2^/sec) because the size of Cy5 is significantly smaller than the Qdot size, it was still lower than most diffusion coefficients of Cy5-labeled APE1, ensuring that APE1 slides along DNA without helical rotation.

**Figure 4.**
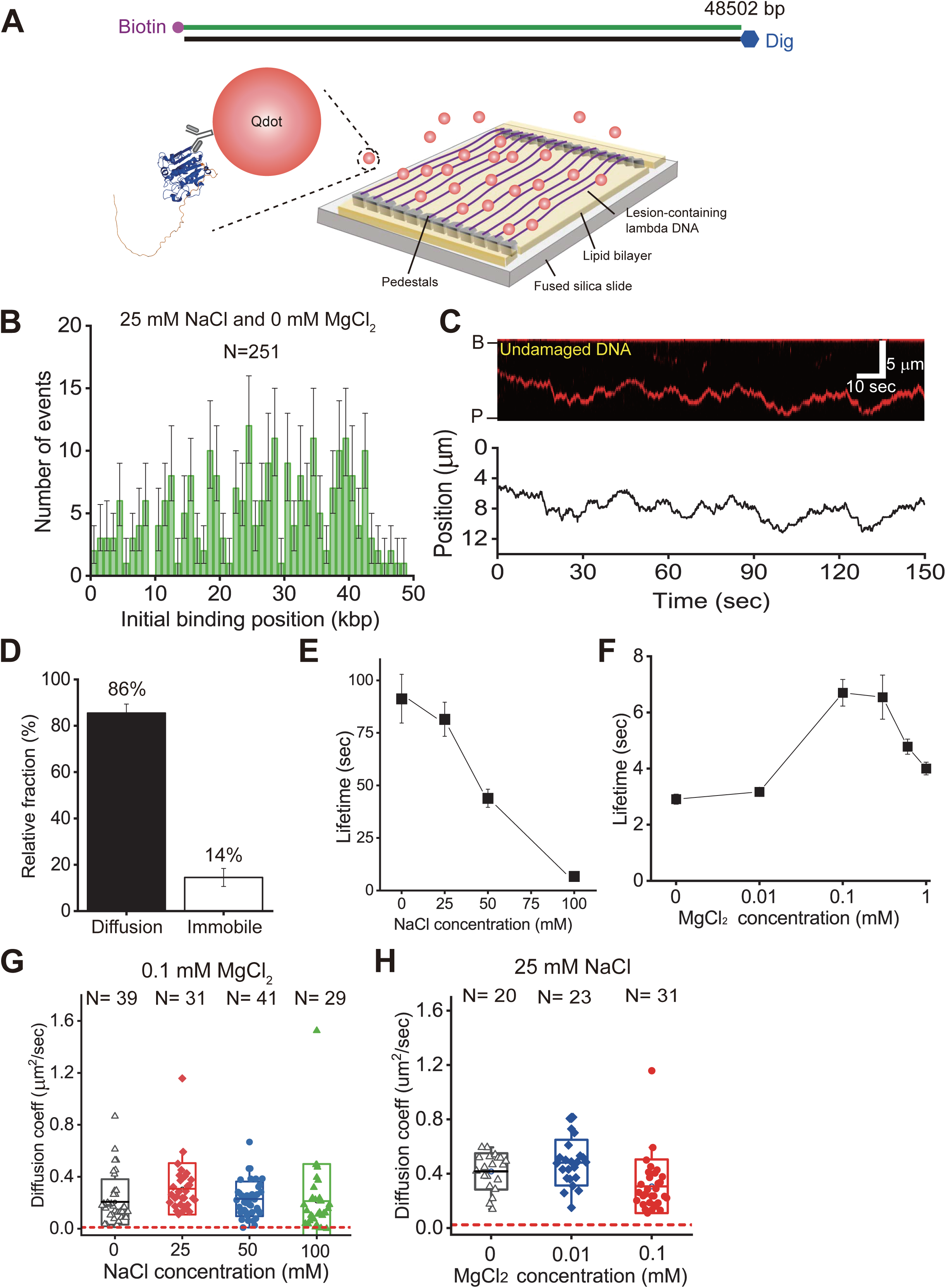
DNA curtain assay of Qdot-labeled APE1. (A) Schematic of the DNA curtain setup. (B) Histogram of APE1 initial binding positions along undamaged λ-DNA. Error bars represent 70% confidence intervals from bootstrap resampling. (C) Kymograph of APE1 diffusion (top) and corresponding particle tracking trace (bottom). (D) Relative fraction of diffusive and immobile APE1 molecules (n = 124). Error bars indicate the standard error of the mean (s.e.m.). (E, F) Diffusion lifetimes at varying NaCl (E) and MgCl₂ (F) concentrations. (G, H) Box plots of diffusion coefficients (D_diff) under varying NaCl (G) and MgCl₂ (H) concentrations. Red dashed lines denote the theoretical limit of D_diff for helical rotation along DNA (0.004 µm²/s). Median ± SD are shown.

### Effect of IDR on the diffusion of APE1 along extended DNA

To investigate the effect of IDR on 1D diffusion along long DNA, the behavior of 0× and 2× IDR mutants was observed using DNA curtains (Figure 3A). Both mutants also exhibited diffusive motion (Figure 5A and 5B). Like wild-type, the diffusion coefficients of 0× IDR mutant were insensitive to ionic strength, indicating that the absence of IDR does not affect the diffusive motion of APE1 (Figure 5C). We next examined the effect of IDR on the diffusion of APE1. As the IDR length increased, the diffusion lifetime was correspondingly extended, suggesting that the interaction between IDR and DNA makes APE1 stay longer on DNA (Figure 5D and Figure S8K-S8M). On the other hand, the length of IDR did not change the diffusion coefficients, suggesting that APE1 diffusion is attributed to the nuclease domain, which maintains the contacts with DNA (Figure 5E). Given that IDR contributes to the diffusion lifetime, IDR-independent diffusion coefficients imply that IDR transiently interacts with DNA to prevent dissociation, consistent with smFRET data.

**Figure 5.**
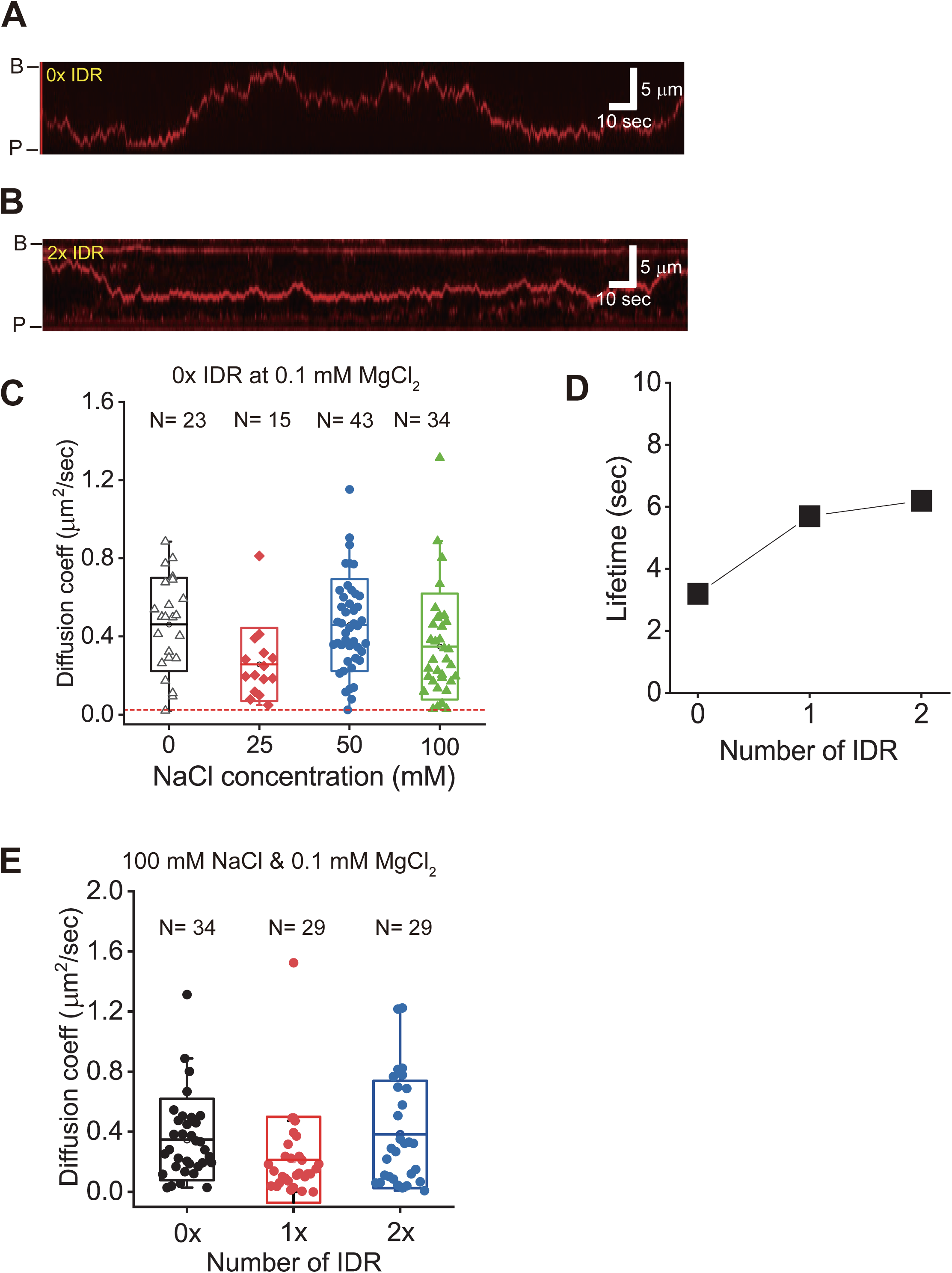
Effect of IDR length on APE1 diffusion. (A, B) Representative kymographs of APE1 lacking IDR (A) and with 2× IDR (B). B and P denote barrier and pedestal, respectively. (C) Box plots of D_diff for 0× IDR APE1 at NaCl concentrations of 0, 25, 50 and 100 mM (median ± SD). (D) Diffusion lifetimes at 100 mM NaCl and 0.1 mM MgCl₂ as a function of IDR length. Error bars represent s.e.m. (E) D_diff values at 100 mM NaCl and 0.1 mM MgCl₂ for APE1 with 0×, 1× and 2× IDR (median ± SD).

### APE1 finds AP sites via 1D diffusion

We next investigated whether APE1 finds AP sites via 1D diffusion using DNA curtain assays with lambda DNA containing AP sites at a specific location (Figure 6A and Figure S7). In DNA curtains, Qdot-labeled APE1 molecules were preferentially bound to the AP sites (Figure 6B and 6C). We also observed that APE1 recognized the AP sites via either direct binding (3D collision) or 1D diffusion (Figure 6D and 6E). APE1 could re-engage with AP sites via 1D diffusion after it dissociated (Figure 6F). The probability of AP site recognition via 1D diffusion was ∼ 66% (Figure 6G). However, this value was underestimated because APE1 could diffuse within our spatial resolution (∼ 1 kbp/pixel). Therefore, our data demonstrated that APE1 dominantly finds AP sites via 1D diffusion, which facilitates the rapid search for AP sites.

**Figure 6.**
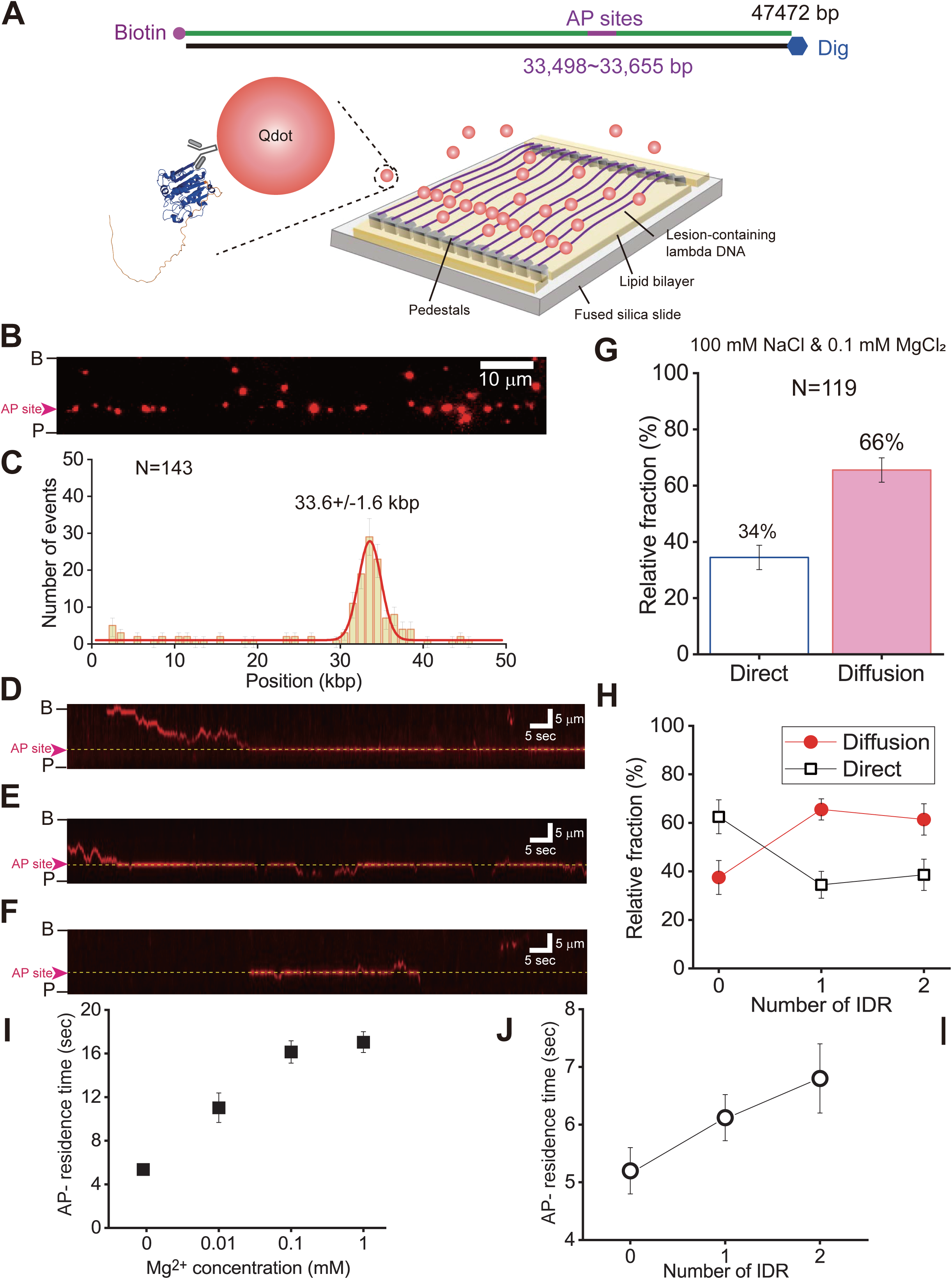
Mechanism of AP site search by APE1 in DNA curtain assays. (A) Schematic of λ-DNA containing three AP sites (33,498–33,655 bp). (B) DNA curtain image showing APE1 binding at AP sites (magenta triangles). (C) Histogram of APE1 binding positions on AP-containing λ-DNA, with a peak at ∼33.6 kb obtained via Gaussian fitting. (D–F) Representative kymographs of APE1 searching for AP sites via 1D diffusion (D), 3D collision (E), and 3D binding followed by 1D diffusion (F). (G, H) Relative fractions of each search mode for wild-type (G) and various IDR lengths (H). Error bars represent standard deviations from binomial distributions. (I) AP site residence times at different MgCl₂ concentrations. (J) Residence times as a function of IDR length. Error bars indicate s.e.m.

Our data above and previous studies suggested that IDR could enhance its interaction with DNA through continual transient contacts with the DNA ^37, 38^. To examine how IDR affects the search for AP sites, we observed the interaction between AP sites and 0× or 2× mutants. 0× mutant dominantly showed 3D collision to find AP sites, whereas wild-type (1×) and 2× mutant had higher fraction for 1D diffusion, suggesting that IDR itself significantly contributes to the diffusive motion of APE1 independently of the length (Figure 6H). We next estimated the AP-residence time meaning by how long APE1 remains on the AP sites. As the Mg^2+^ concentration increased, AP-residence time of wild-type at the AP sites increased (Figure 6I and Figure S9A-S9D). Structurally, Mg^2+^ is coordinated near the DNA binding pocket of APE1 by a cluster of negatively charged residues, including D70, E96, and D308 ^39, 40^. This local electrostatic environment likely generates repulsion against the DNA phosphate backbone. Our data suggest that Mg²⁺ neutralizes these repulsive charges, stabilizing the interaction of APE1 with DNA and prolonging its residence time at AP sites. Given that IDR contributes to binding with undamaged DNA (Figure S5), it is likely that the IDR mediates transient electrostatic interaction with the DNA backbone, thereby facilitating initial engagement and promoting stabilization at lesion sites. Consistently, the residence time of APE1 at AP sites increased with IDR length, indicating that IDR enhances the stability of APE1-DNA interaction at sites of damage, e.g., AP sites (Figure 6J and Figure S9E-S9G).

### Catalytic dead mutant D210N behaves similarly to wild-type

To accurately monitor DNA search dynamics of APE1 without the confounding effect of DNA cleavage, we utilized a catalytically inactive mutant (D210N), in which the aspartate residue at position 210—critical for coordinating a water molecule during phosphodiester bond cleavage ^39^—is substituted with asparagine. Structural studies have shown that D210, along with E96, D70, and D308, participates in coordinating a single Mg²⁺ ion at the active site, stabilizing the catalytic center (Figure 2A). The D210N mutant disrupts this coordination, abolishing cleavage activity (Figure S2A) while preserving the structural integrity of the binding interface (Figure S2B and S2C). In our DNA curtain assays conducted at 0.1 mM Mg²⁺, we confirmed that D210N retains AP site recognition capability and exhibits DNA-binding behavior comparable to wild-type APE1 (Figure S10). Specifically, D210N showed similar frequencies of 1D diffusion and 3D collision for AP site targeting (Figure S10E), as well as comparable diffusion coefficients and diffusion lifetimes (Figure S10F–G), indicating that cleavage activity is dispensable for lesion search. Notably, the AP-residence time of D210N was approximately 2.5 times increased compared with that of wild-type presumably because D210N stays at AP sites without cleaving DNA backbone (Figure S10H and Figure S9H). At an AP site, D210 coordinates with a water molecule for interacting with phosphate backbone ^39, 41^. In D210N, the charge neutralization due to glutamine disturbs the nucleophilic attack of water molecule to DNA backbone. It also would reduce the electrostatic repulsive interaction, and hence the AP-residence time of D210N could be increased. Together, these observations validate the use of D210N as a non-cleaving surrogate for wild-type APE1 in DNA curtain assays, particularly under conditions where Mg²⁺ is required to support physiological diffusion dynamics.

### Effect of IDR on the diffusion of APE1 in molecular dynamics simulations

To investigate the diffusion mechanism of APE1 and the mechanistic roles of its IDR, we performed atomistic molecular dynamics (MD) simulations of wild-type APE1 and the 0× IDR mutant in the vicinity of DNA (Figure 7A). Our MD simulation framework is particularly well-suited to capture dynamic behaviors such as transient and nonspecific protein–DNA interactions ^42^, thereby providing mechanistic insights into how the IDR modulates 1D diffusion of APE1 along DNA. In contrast, structural models of the APE1–DNA complex generated using AlphaFold3 ^43^ predict the N-terminal IDR in an extended conformation without any significant contacts with DNA (Figure 7B). This highlights a limitation of current AI-based structure prediction algorithms in capturing the functional roles of IDRs, especially in processes involving diffusion.

**Figure 7.**
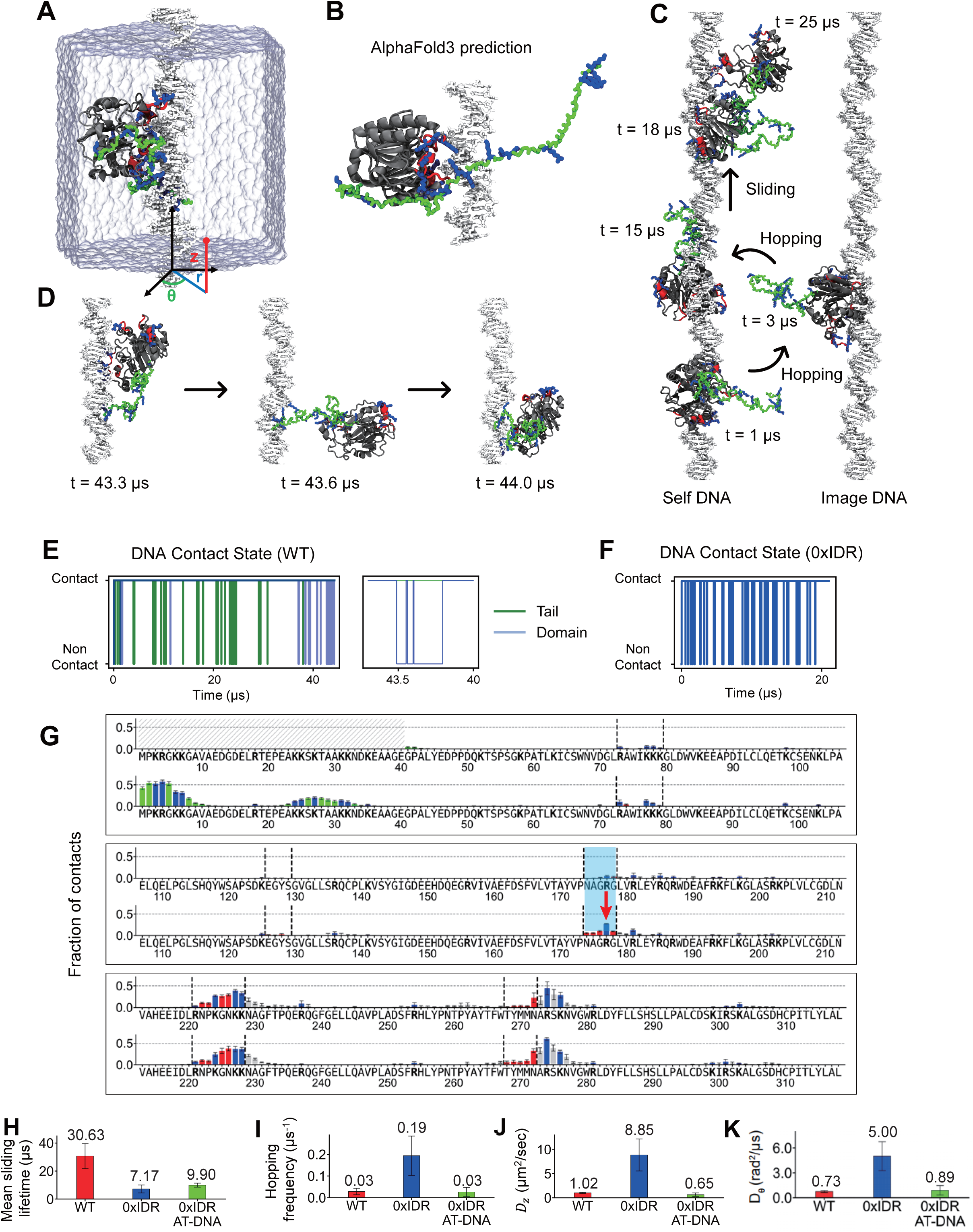
Molecular dynamics (MD) simulations reveal IDR-dependent modulation of APE1 diffusion along DNA. (A) Initial configuration of the MD simulation system showing wild-type APE1 placed near a periodic DNA duplex aligned along the z-axis. Canonical DNA-binding residues in the crystal structure are shown in red; lysine (K) or arginine (R) residues are highlighted in blue ^17^. The inset illustrates the cylindrical coordinate system used to quantify translational (*z*) and rotational (*θ*) diffusion. (B) AlphaFold3-predicted model of the APE1–DNA complex ^43^. (C) Representative snapshots from the simulations of wild-type APE1 showing sliding along DNA and hopping between periodic image helices. (D) Time series of a transient dissociation event followed by rapid rebinding to the original self-DNA, facilitated by anchoring of the IDR. (E) Time-dependent DNA contact analysis for the structured domain and IDR tail in the wild-type APE1 system. The blue and green lines denote the domain-DNA and tail-DNA contacts, respectively. The right panel shows the interval from 43 µs to 44 µs. (F) Time-dependent DNA contact analysis for the structured nuclease domain in the mutant APE1 system. (G) Residue-wise DNA contact fraction averaged over all four simulations of wild-type APE1, shown for the 0× IDR mutant (top) and the 1× IDR WT (bottom). Canonical DNA-binding residues identified in the crystal structure are marked in red; K/R residues are marked in blue ^17^. Remarkably, acquisition of the IDR enables persistent interactions mediated by an unprecedented 177 arginine residue (red arrow in the sky-blue window) within the structured catalytic domain, prolonging residence time and facilitating long-range diffusion. (H–J) Quantitative comparison of sliding lifetime (H), hopping frequency (I), axial translational diffusion coefficient (J), and rotational diffusion coefficient (K) for wild-type (red), 0× IDR mutant (blue), and 0× IDR mutant on AT-rich DNA (green).

All simulations employed periodic boundary conditions, with the DNA duplex aligned along the z-axis (Figure 7A). The DNA termini were covalently bonded and positioned to ensure seamless continuity across the periodic boundaries, effectively rendering both DNA strands infinite in length ^44^. By eliminating end effects, this configuration offers a setting ideally suited for analyzing diffusion along DNA ^42^. APE1 was initially placed at random positions and orientations relative to the DNA in each simulation (Figure 7A). For both the wild-type APE1 and the 0× IDR mutant, we carried out four independent replicates to ensure statistical robustness. Simulations were performed using the AMBER ff99SB-ILDN-phi force field ^45–47^ with the CUFIX corrections ^42, 48^ and the GROMACS package ^49^. See the Methods section for simulation details.

Simulation trajectories of wild-type APE1 revealed that both the IDR and the structured nuclease domain engage in transient yet persistent interactions with DNA, collectively supporting translational and rotational diffusion along the helical axis (Figure 7C and Supplementary Movies S1–S4). Despite the highly dynamic nature of this 1D diffusion process, the orientation of APE1 relative to the DNA remained closely aligned with that observed in the crystal structure ^50^, suggesting that the protein retains a functionally relevant binding pose during 1D diffusion (Figure 7C). To further characterize these interactions, we quantified the residue-wise DNA contact fraction averaged over the entire simulation (Figures 7E and S11). This analysis revealed that residues constituting the canonical DNA-binding interface in the crystal structure ^50^ predominantly remain in contact with DNA during diffusion, exhibiting mean contact fractions as high as 0.87. Notably, the N-terminal segment of the IDR with multiple lysine or arginine residues (residues 3–7 and 24–27) displayed a high mean contact fraction of 0.65, highlighting its sustained binding to DNA during 1D diffusion (Figure 7C).

Interestingly, in addition to continuous diffusion along DNA, we observed transient dissociation events of the nuclease domain, during which the wild-type APE1 briefly dissociates from the DNA before rebinding to either the same DNA helix (self-DNA) (Figure 7D) or hopping to a neighboring periodic image DNA across the periodic boundary (Figures 7C and S12A). Rebinding to self-DNA frequently coincided with the IDR remaining anchored to the same DNA molecule (see the center of Figure 7D), suggesting that the IDR facilitates rapid reassociation and thereby suppresses long-range hopping or complete dissociation. Exemplary time-dependent DNA contact analyses showed that APE1 with one IDR copy (1× IDR) maintained longer and more stable DNA contact time (i.e., dwell time on the top of Figure 7E) than the IDR-deleted mutant (0× IDR; Figure 7F). Moreover, residue-wise DNA contact fractions averaged over simulations (Figure 7G) revealed that acquisition of the IDR in the 1× IDR WT APE1 (bottom), compared with the 0× IDR mutant (top), confers persistent interactions mediated by a previously unrecognized 177 arginine residue (red arrow in the sky-blue inset) within the structured nuclease domain, thereby prolonging residence time and enabling long-range diffusion. Consistently, deletion of the IDR markedly reduced the mean sliding lifetime, decreasing—from 31 μs in the wild-type to 7 μs in the 0× IDR mutant (Figure 7H)—while simulations of 0× IDR exhibited substantially more frequent hopping events between periodic image DNA helices (Figures 7I and S13A), in agreement with our experiments (Figure 3E and 5D). Specifically, five hopping events occurred over a cumulative 185 μs of simulation for wild-type APE1 (Figure S12), whereas sixteen hopping events were detected within 70 μs for the 0× IDR mutant (Figures 7H and S13).

To quantitatively assess the diffusion modes of APE1, we analyzed the motion of the center of mass (CoM) of the structured domain in a cylindrical coordinate system fixed to the DNA helix (Figure 7A). Specifically, we tracked the axial position (*z*) and angular orientation (*θ*) of the CoM as functions of time (Figures S12, and S13). In this *z* − *θ* space, APE1 exhibited periods of random diffusion interspersed with intervals of helical motion that appeared to follow the DNA groove structure. To distinguish these two modes, we computed the translational diffusion coefficient along the DNA axis (*D*_*z*_, in µm^2^/sec) and the rotational diffusion coefficient around the DNA axis (*D*_*θ*_, in rad^2^/µsec). Under perfect groove tracking, translational and rotational motion are geometrically coupled by the DNA helical pitch *P* = 3.4 nm, yielding the relation: 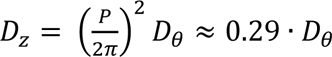. Thus, the ratio 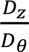 serves as a quantitative metric to distinguish between groove tracking and random diffusion modes.

For wild-type APE1, we measured *D*_*z*_ ≈ 1.02 µm²/sec and *D*_*θ*_ ≈ 0.73 rad^2^/µsec (red bars in Figures 7J and 7K), yielding a ratio 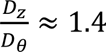. This significantly exceeds the theoretical prediction for perfect groove tracking (∼0.29), indicating that APE1 translates along the axis more extensively than expected for strict groove tracking. These results suggest partial decoupling between rotational and translational motion, with APE1 alternating between groove-tracking and more linear sliding modes. This observation is consistent with our DNA curtain assays, which indicate that APE1 predominantly slides along the DNA axis without significant rotation around the DNA axis.

For wild-type APE1, the simulated *D*_*z*_was ∼1.0 µm²/sec (Figure 7J), modestly overestimating the experimentally measured value of ∼0.4 µm²/sec (Figure 5C and 5G). This discrepancy is reasonable given that the DNA in simulations is modeled as perfectly straight, whereas the DNA in DNA curtain assays often exhibits curvature. In contrast, the 0× IDR mutant showed *D*_*z*_∼8.9 µm²/sec in simulations, markedly higher that the experimental *D*_*z*_ (Figure 7J). We attribute this discrepancy, at least in part, to the use of a GC-rich DNA sequence, which has been shown to reduce protein–DNA attraction relative to AT-rich or mixed sequences ^51^. Supporting this interpretation, additional simulations of the 0× IDR mutant using AT-rich DNA yielded a much lower *D*_*z*_ ∼0.7 µm²/sec, suggesting that in more physiologically relevant, mixed-sequence contexts, *D*_*z*_ would more match experimental measurements.

## Discussion

Our single-molecule analyses reveal that APE1 predominantly recognizes AP sites through 1D diffusion rather than stochastic 3D collisions. This suggests a guided scanning mechanism, wherein APE1 diffuses along DNA duplex to locate lesions. Notably, the diffusion coefficients were largely independent of ionic strength, supporting a sliding mechanism over hopping. This behavior is functionally advantageous, as the sliding mode enables APE1 to sense local DNA flexibility, a biophysical hallmark of AP sites. AP-containing DNA is thermodynamically destabilized and often exhibits 20–30° kinks relative to regular duplex DNA ^52–54^. Coupled with the known structural observation that APE1 bends DNA by approximately −35° to accommodate an AP site into its catalytic pocket ^55^, these findings support a model in which APE1 exploits structural distortions in DNA to locate its substrates efficiently.

To further dissect the factors influencing APE1’s diffusion dynamics, we examined the role of Mg²⁺, a divalent metal ion known to be essential for its catalytic activity. In addition to promoting phosphodiester bond cleavage at AP sites, Mg²⁺ markedly enhanced APE1’s ability to slide along DNA. Physiological concentrations of Mg²⁺ increased both diffusion lifetimes and translocation frequencies during 1D scanning (Figures 2B–C and 4F). By contrast, mutation of the conserved glutamate residue E96, which directly coordinates Mg²⁺ in the active site, abolished this enhancement (Figure 2D), highlighting the importance of Mg²⁺-mediated electrostatic stabilization for APE1–DNA interactions. Structural analyses further support this model. In the absence of Mg²⁺, the negatively charged residues at the active site—E96, D70, and D308—are likely to experience electrostatic repulsion from the DNA phosphate backbone, destabilizing APE1’s interaction with DNA during diffusion. When Mg²⁺ is coordinated at the catalytic center, it neutralizes these local negative charges, thereby reducing repulsion and strengthening APE1’s transient contact with the DNA. This allows for longer diffusion lifetimes and more sustained 1D scanning, even on undamaged DNA.

While Mg²⁺ facilitates diffusion through charge shielding at the active site, APE1 also possesses an N-terminal intrinsically disordered region (IDR) that has been implicated in multiple nuclear functions, including redox regulation of transcription factors such as p53, NF-κB, and HIF-1α, independently of its DNA repair function ^56^. More recently, the IDR has been linked to nucleolar localization and stress-responsive condensate formation, mediated through interactions with RNA and stress signaling proteins ^57–59^. However, these prior studies have primarily focused on the ensemble-level functions of the IDR. Whether and how the IDR contributes to APE1’s lesion search dynamics at the single-molecule level has remained unexplored. To address this gap, we used smFRET and DNA curtain assays to directly probe the role of the IDR in APE1’s DNA scanning behavior. Deletion of the IDR significantly reduced the frequency of translocation events between two AP sites (Figure 3B), indicating that the IDR facilitates efficient 1D sliding (Figure 3C). This result was corroborated by DNA curtain experiments, where the IDR-deficient mutant showed shorter diffusion lifetimes and reduced residence time at AP sites (Figures 5D and 6J). Moreover, loss of the IDR shifted APE1’s lesion search mechanism toward 3D-based sampling, further underscoring its importance in enabling efficient linear diffusion (Figure 6H).

To determine whether the IDR’s effect was length-dependent, we generated an IDR-duplicated mutant and observed a substantial enhancement in APE1’s diffusion characteristics. APE1 with duplicated IDR exhibited prolonged scanning lifetimes and increased translocation frequency compared to the wild-type protein (Figures 3C and 5E), suggesting that the IDR supports diffusion in a scalable manner. Besides, a cooperative mechanism wherein Mg²⁺ coordination at the catalytic site and the IDR at the N-terminus work in concert to promote 1D diffusion of APE1. This dual strategy allows APE1 to maintain a balance between dynamic scanning and stable target engagement, enabling robust and efficient lesion recognition within the chromatin landscape. This integrated mechanism may reflect a general principle used by eukaryotic DNA repair enzymes to navigate the genome and maintain DNA integrity under physiological conditions.

MD simulations further support our experimental findings. Atomistic MD simulations of wild-type APE1 and the IDR-deficient mutant revealed that both the nuclease domain and the IDR establish transient yet persistent contacts with DNA, facilitating a combination of translational and rotational sliding. Regarding the role of disordered and structured regions of APE1 in 1D diffusion, contact probability analysis (Figure 7E) revealed that the N-terminal arginine and lysine residues—especially those within residues 1–30—displayed significant DNA-binding frequencies (contact fraction > 0.6), highlighting their key role in anchoring the IDR to the DNA helix. These positively charged residues likely form electrostatic bridges with the negatively charged phosphate backbone, stabilizing APE1 on the DNA during scanning. In addition to the IDR, the structured nuclease domain of APE1 also contributed significantly to DNA binding during diffusion. Contact mapping revealed that a cluster of lysine and arginine residues near the catalytic core—particularly those corresponding to canonical DNA-binding regions (Figure 7E, red and blue bars)—remained engaged with the DNA duplex during sliding. These basic residues likely help to maintain proper orientation and alignment of the endonuclease domain with the DNA helix, ensuring readiness for lesion recognition and catalysis. Such contacts are critical not only for the stability of the scanning complex but also for positioning the active site for efficient engagement of lesions of AP sites.

## Data availability

The smFRET and DNA curtain data are included as supplementary and source data with the paper. Simulation trajectories are deposited at https://zenodo.org/records/15833711. Source data are provided with this paper.

## Supplementary data

Supplementary data is available at NAR online.

## Acknowledgements

We thank all members of the J.Y., J.Y.L, and G.L. labs for assistance and helpful discussion. This research was supported by KAIST Grand Challenge 30 Project (KC30) and grants from the National Research Foundation of the Korea (RS-2024-00408712 and RS-2024-00341654) and Korea Drug Development Fund (RS-2024-00463605). This research was supported by the National Research Foundation of Korea grants to J.Y.L. (grant number RS2024-00449988) and the Institute for Basic Science (grant number IBS-R022-D1). J.Y. and G.J. were supported by the National Research Foundation of Korea (NRF) grant (No. 2020R1A2C1101424) and Institute of Information & communications Technology Planning & Evaluation (IITP) grant (No.RS-2023-00220628).

## Author contributions

J.Y., J.Y.L, and G.L. conceived the project. D.L., J.K., and J.Y. performed the single-molecule experiments; S.K. conducted DNA curtain assays, biochemical assays and their analyses; and J.Y. and G.J. jointly designed, conducted, and analyzed the MD simulations. All authors wrote the paper.

## Competing interests

The authors declare no competing interests.

## Notes

### Competing Interest Statement

The authors have declared no competing interest.

