## Supplementary Information for "APE1 Coordinates Its Disordered Region and Metal Cofactors to Drive Genome Surveillance"

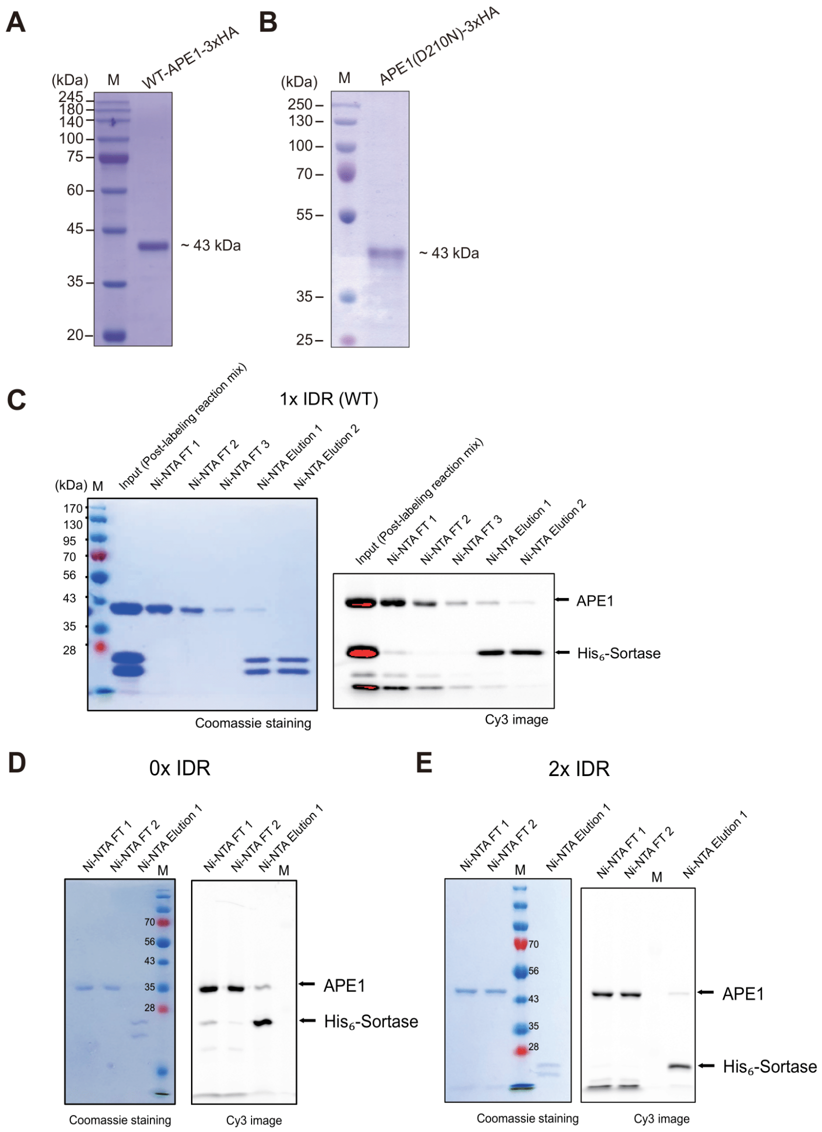


**Supplementary Figure 1. Purification of APE1 constructs and mutants and Sortase-mediated N-terminal Cy3 labeling.** SDS PAGE for purified (A) wild-type-3xHA, (B) D210N-3xHA, (C) wild-type APE1 (1 × IDR), (D) 0 × IDR APE1, and (E) 2 × IDR APE1. For each construct, the left sub-panel is a Coomassie-stained 12 % SDS-PAGE image, and the right sub-panel is the corresponding Cy3 fluorescence image of the same gel. Lanes : M, molecular-weight marker; Input, post-labeling reaction mixture containing Cy3-APE1 and $His_6$-sortase; FT 1–3, Ni-NTA flow-through fractions that recover Cy3-labeled APE1 lacking a His tag; Elution 1–2, 250 mM imidazole elution fractions enriched in $His_6$-Sortase (~20 kDa). Coomassie staining confirms comparable protein loads and shows that Sortase is largely confined to the elution fractions, whereas Cy3 fluorescence appears exclusively at the APE1 bands (≈ 38 kDa for 1 × IDR, ≈ 33 kDa for 0 × IDR, and ≈ 43 kDa for 2 × IDR) with minimal signal in the Sortase band. Together, the images demonstrate efficient N-terminal Cy3 labeling of all three APE1 variants and effective removal of the Sortase enzyme.

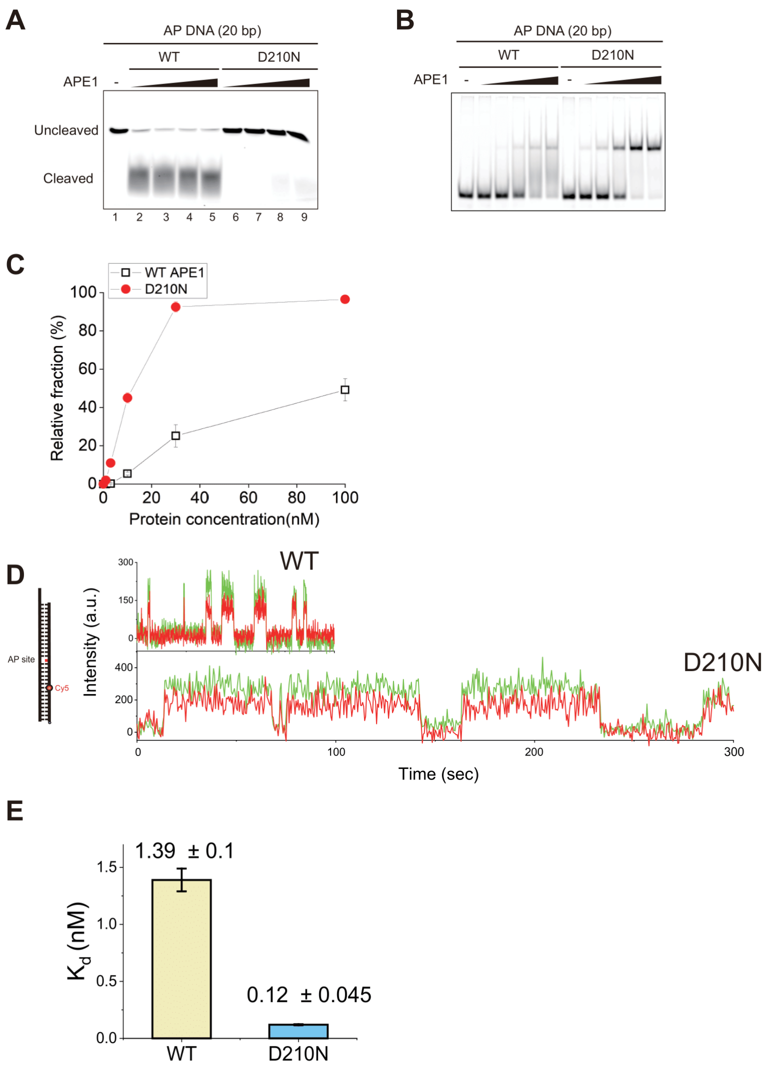


**Supplementary Figure 2. Single-molecule fluorescence analysis reveals that the catalytic-dead APE1 mutant (D210N) binds AP-DNA with higher affinity than wild-type APE1.** (A) Activity test of purified wild-type APE1 and D210N by cleavage of AP-site containing DNA in the presence of 0.1 mM $MgCl_2$. (B) Electrophoretic mobility shift assay (EMSA) for purified wild-type APE1 and D210N in the absence of $MgCl_2$. (C) Quantification of EMSA in B. Error bars are obtained from s.e.m. in triplicate. (D) Representative single-molecule fluorescence time traces. AP-DNA labeled with Cy5 was tested with either wild-type APE1 (top) or the catalytically inactive D210N mutant (bottom) labeled with Cy3. Traces show donor (Cy3, green) and acceptor (Cy5, red) fluorescence intensities over time. Before binding, no fluorescence signal is observed; upon DNA binding, both Cy3 and Cy5 signals appear simultaneously, indicating protein-DNA interaction (E) Dissociation constant ($K_d$) for wild-type APE1 and D210N. $K_d$ values were derived from the ratio $k_{off}/k_{on}$, where $k_{on}$ and $k_{off}$ were obtained by analyzing individual traces per construct (hidden Markov modeling; 100 msec for wild-type APE1 or 500 msec for D210N time resolution). The D210N mutant displays a significantly lower $K_d$, indicating tighter binding to AP-DNA.

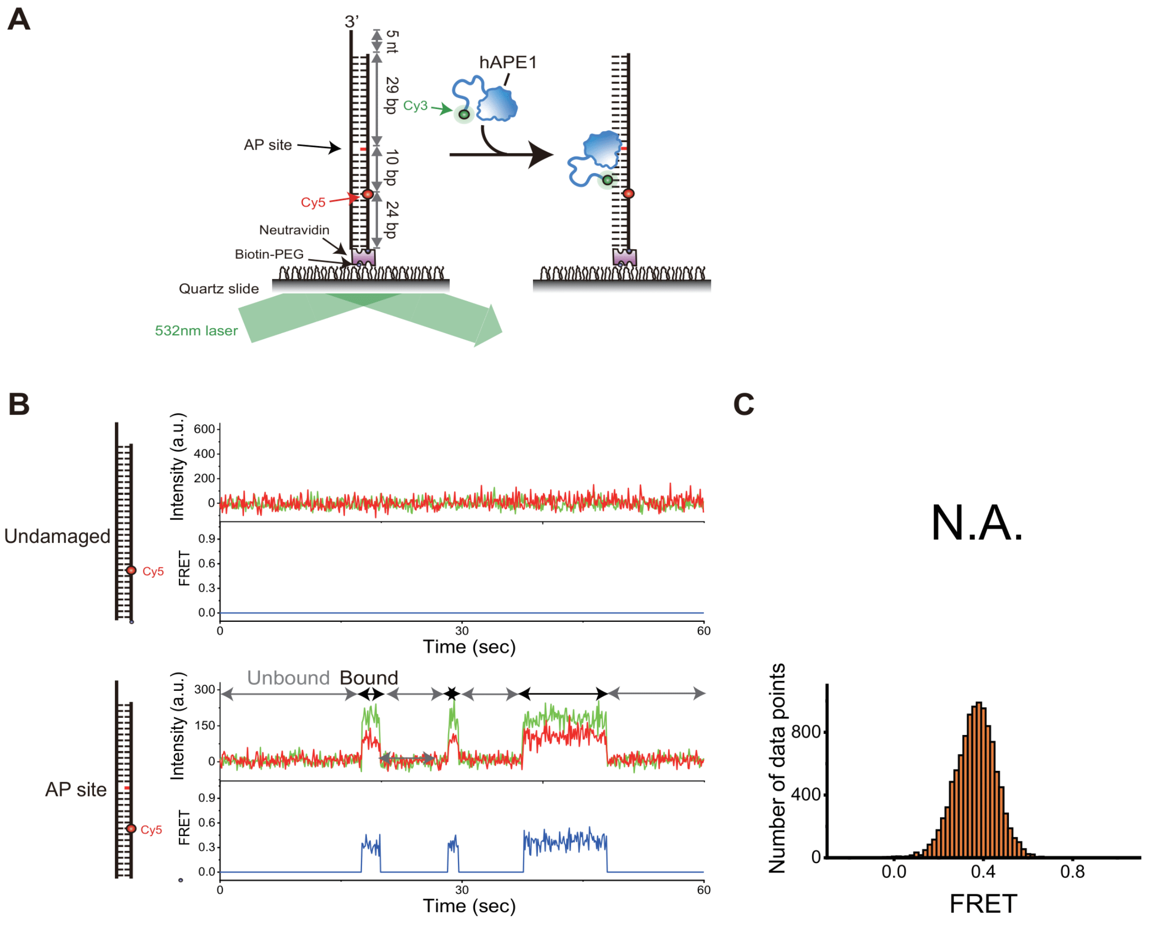


**Supplementary Figure 3. smFRET assay for AP site-specific binding of APE1.** (A) Schematic of the smFRET assay depicting the states before binding (left) and after binding (right). (B) Representative FRET-time trajectories showing APE1 binding events on undamaged DNA and AP-DNA substrates. (C) FRET histograms of APE1 binding events for undamaged DNA, AP-DNA substrates.

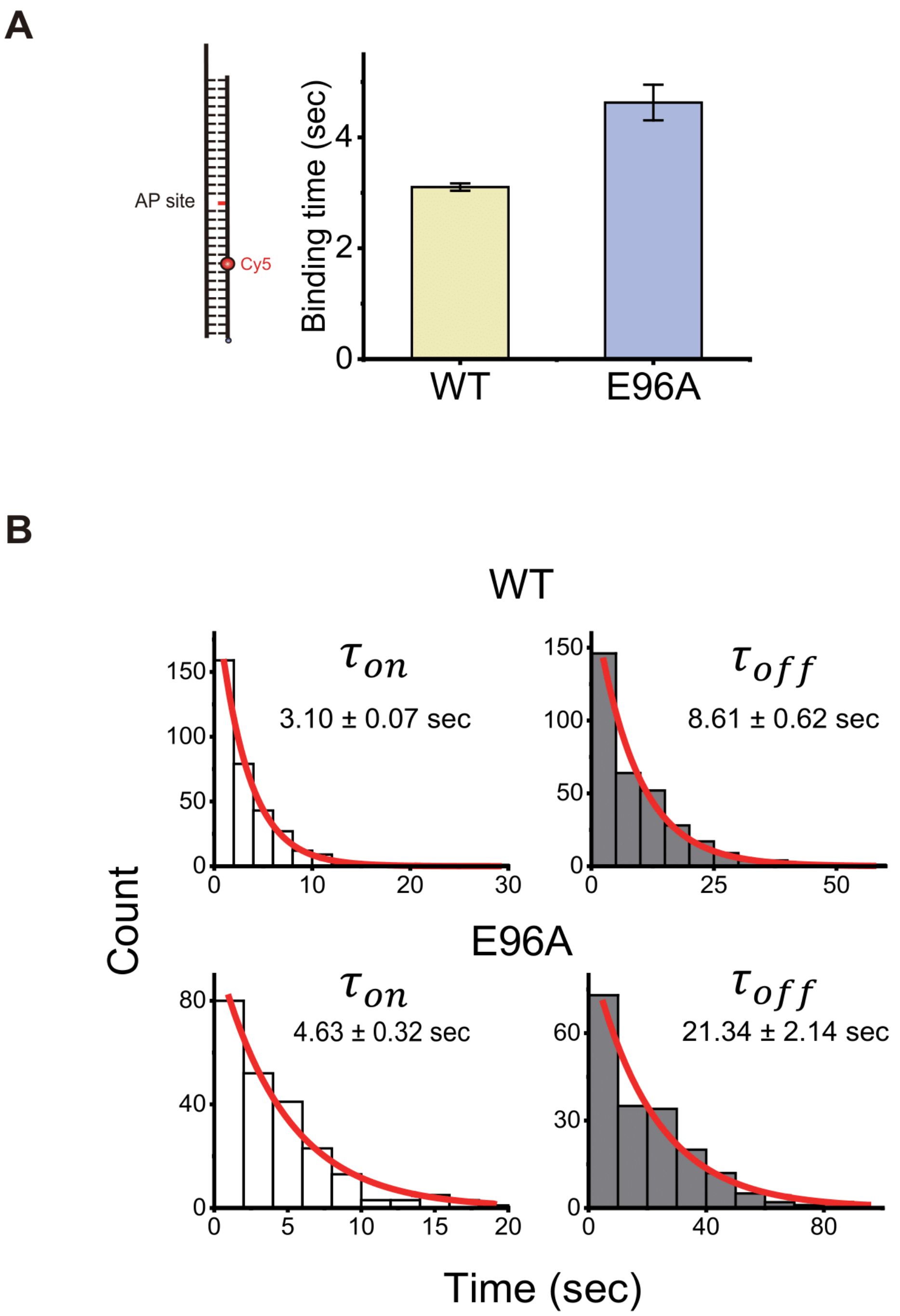


**Supplementary Figure 4. Comparison of AP site binding times between APE1 wild-type and E96A mutant.** (A) Bar graph comparing binding times for wild-type (3.10 ± 0.07 s) and E96A (4.63 ± 0.32 s). (B) Histograms of on times (left) and off times (right) for APE1 wild-type and E96A mutant.

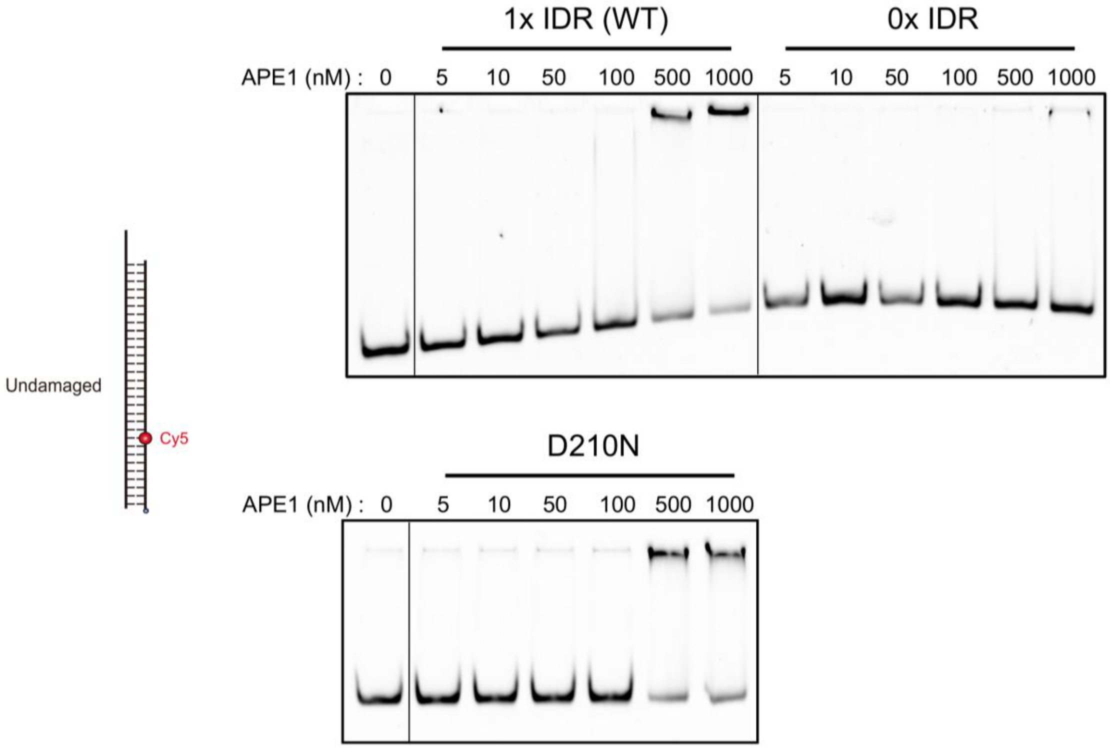


**Supplementary Figure 5. IDR-mediated non-specific binding to undamaged DNA.** EMSA showing the binding of wild-type, 0× IDR and D210N APE1 to undamaged DNA at enzyme concentrations ranging from 0 to 1000 nM.

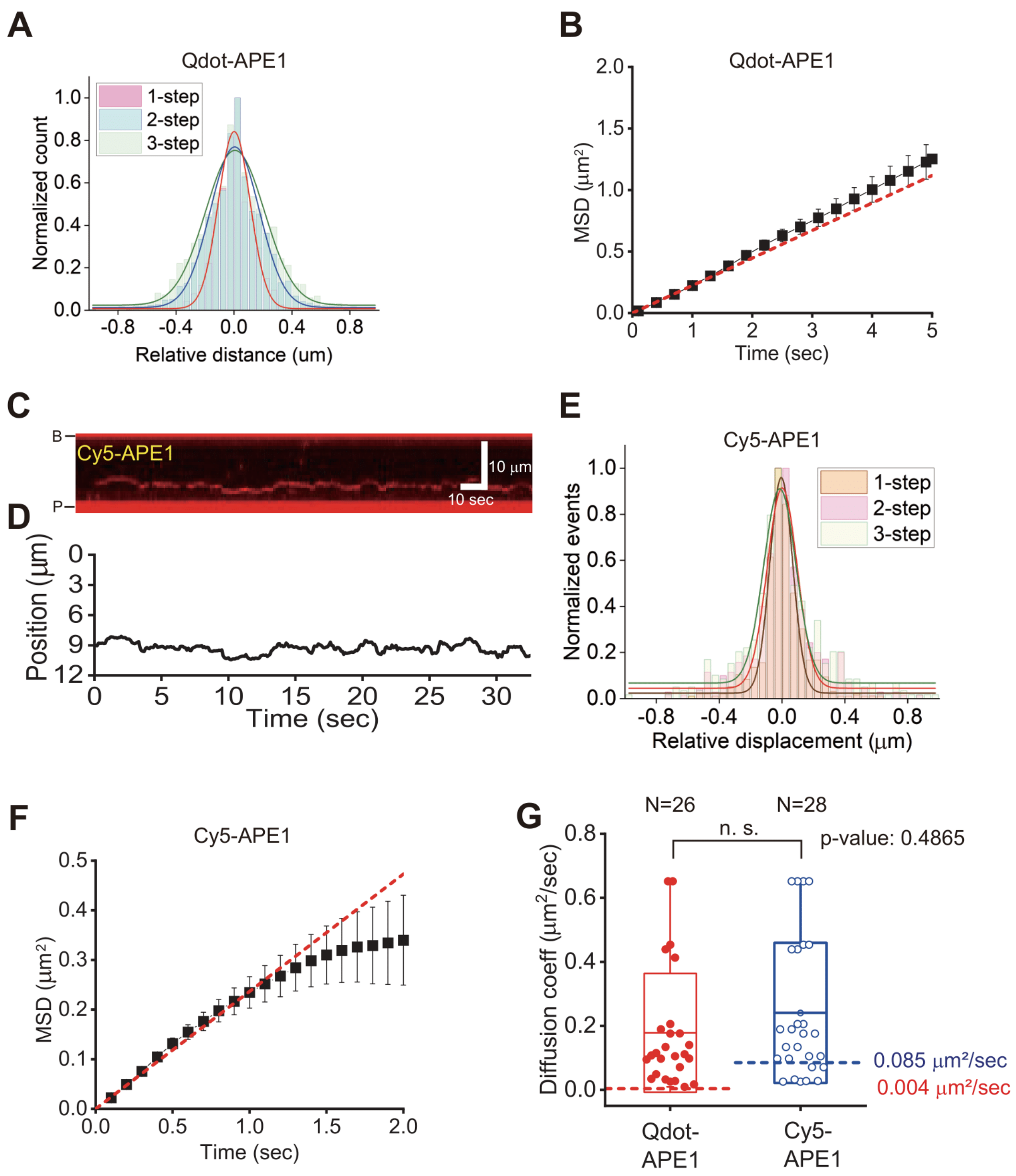


**Supplementary Figure 6. Diffusion analysis of APE1.** (A) Histograms for relative displacements of APE1. (B) Mean square displacement (MSD) of a APE1 molecule. The diffusion coefficient ($D_{diff}$) was estimated from the linear fitting with the first three points of MSD (red dashed line). (C) A representative kymograph for the diffusion of [Cy5]-APE1. (D) Time trace of that was obtained by particle tracking of the molecule in Figure S6C. (E) Histograms for relative displacements of [Cy5]-APE1. (F) MSD of a [Cy5]-APE1 molecule. $D_{diff}$ was estimated from the linear fitting with the first three points of MSD (red dashed line). (G) Comparison of $D_{diff}$ between Qdot-APE1 and [Cy5]-APE1. Both experiments were carried at 100 mM NaCl and 0.1 mM $MgCl_2$. Red and blue dashed lines represent the theoretical limits of diffusion coefficient for helical rotation around DNA.

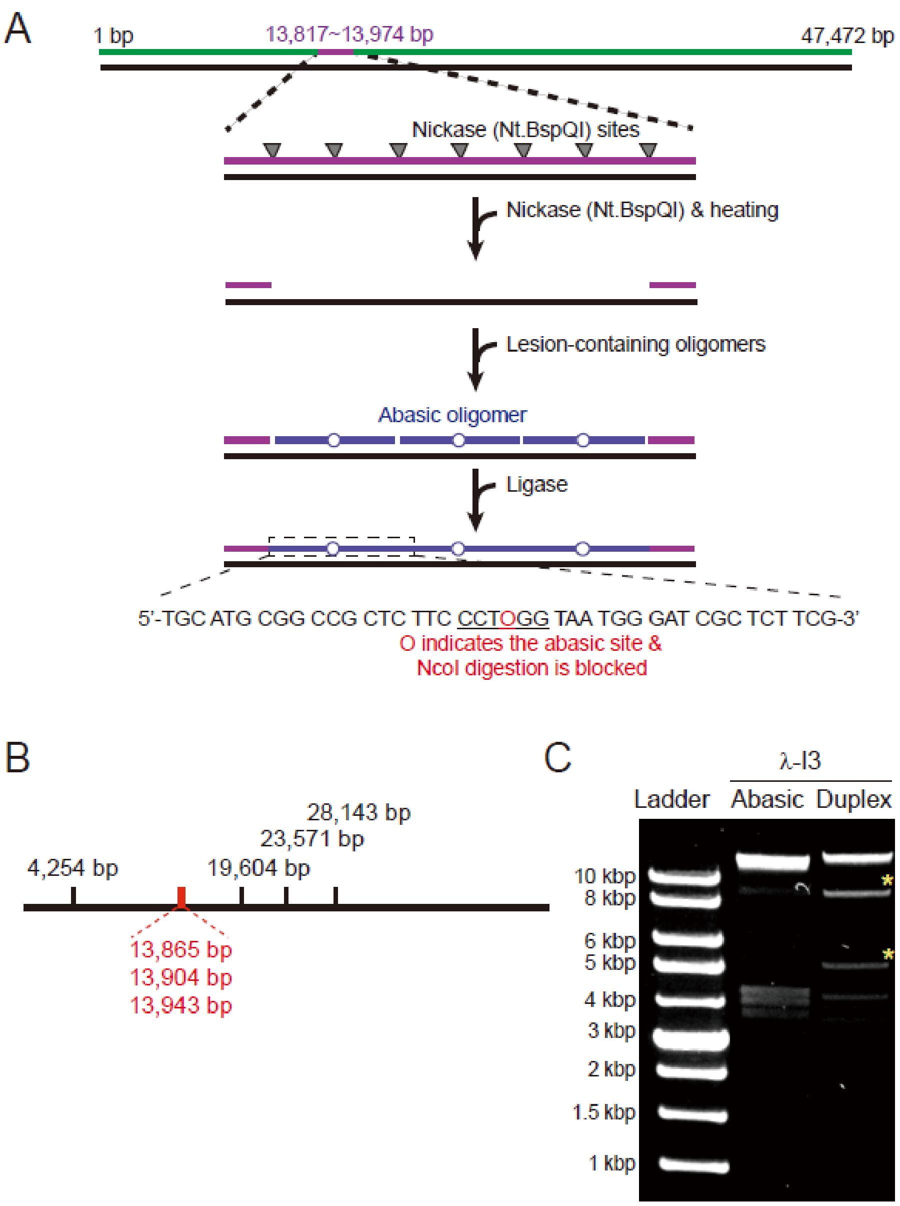


**Supplementary Figure 7. Preparation of lambda DNA containing AP sites.** (A) Insertion of AP sites in engineered lambda DNA (λ-I3), which contains seven nickase sites between 33,514 bp and 33,630 bp. The detailed protocol was described at Methods. (B) Location of NcoI digestion sites in λ-I3. NcoI positions within the seven nickase sites are denoted in red. (C) Agarose gel electrophoresis for NcoI-treated λ-I3 to check the insertion of AP-containing oligomers. When AP oligomers are properly inserted, 9,611 bp and 5,661 bp fragments (yellow asterisks) are not shown in the gel.

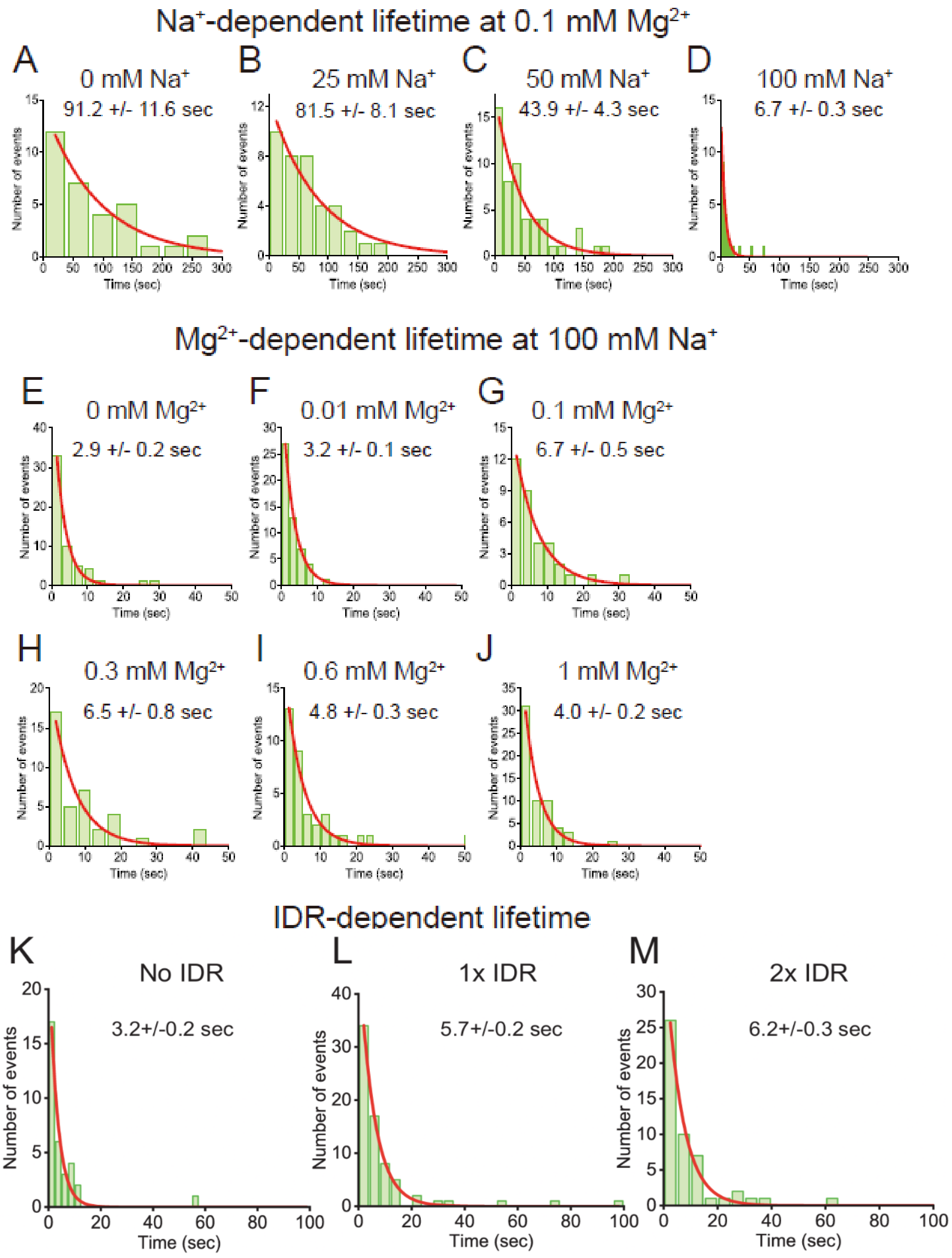


**Supplementary Figure 8. Analyses for diffusion lifetime.** (A-D) Analyses for diffusion lifetime according to NaCl concentration at 0.1 mM $MgCl_2$ in Figure 4E. (E-J) Analyses for diffusion lifetime according to $MgCl_2$ concentration at 100 mM NaCl in Figure 4F. (K-M) Analyses for diffusion lifetime according to the IDR length in Figure 5D.

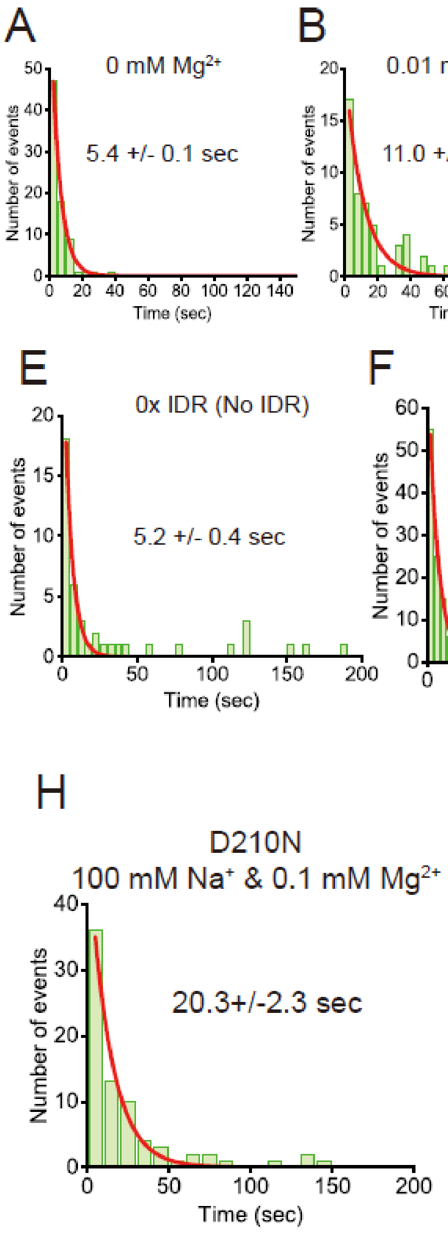


**Supplementary Figure 9. Analyses for abasic residence time.** (A-D) Analyses for abasic residence time according to $MgCl_2$ concentration at 100 mM NaCl in Figure 6I. (E-G) Analyses for abasic residence time according to the IDR length at 100 mM NaCl and 0.1 mM $MgCl_2$ in Figure 6J. (H) Analysis for abasic residence time for D210N at 100 mM NaCl and 0.1 mM $MgCl_2$ in Figure 7H. All graphs are fitted with a single exponential decay function.

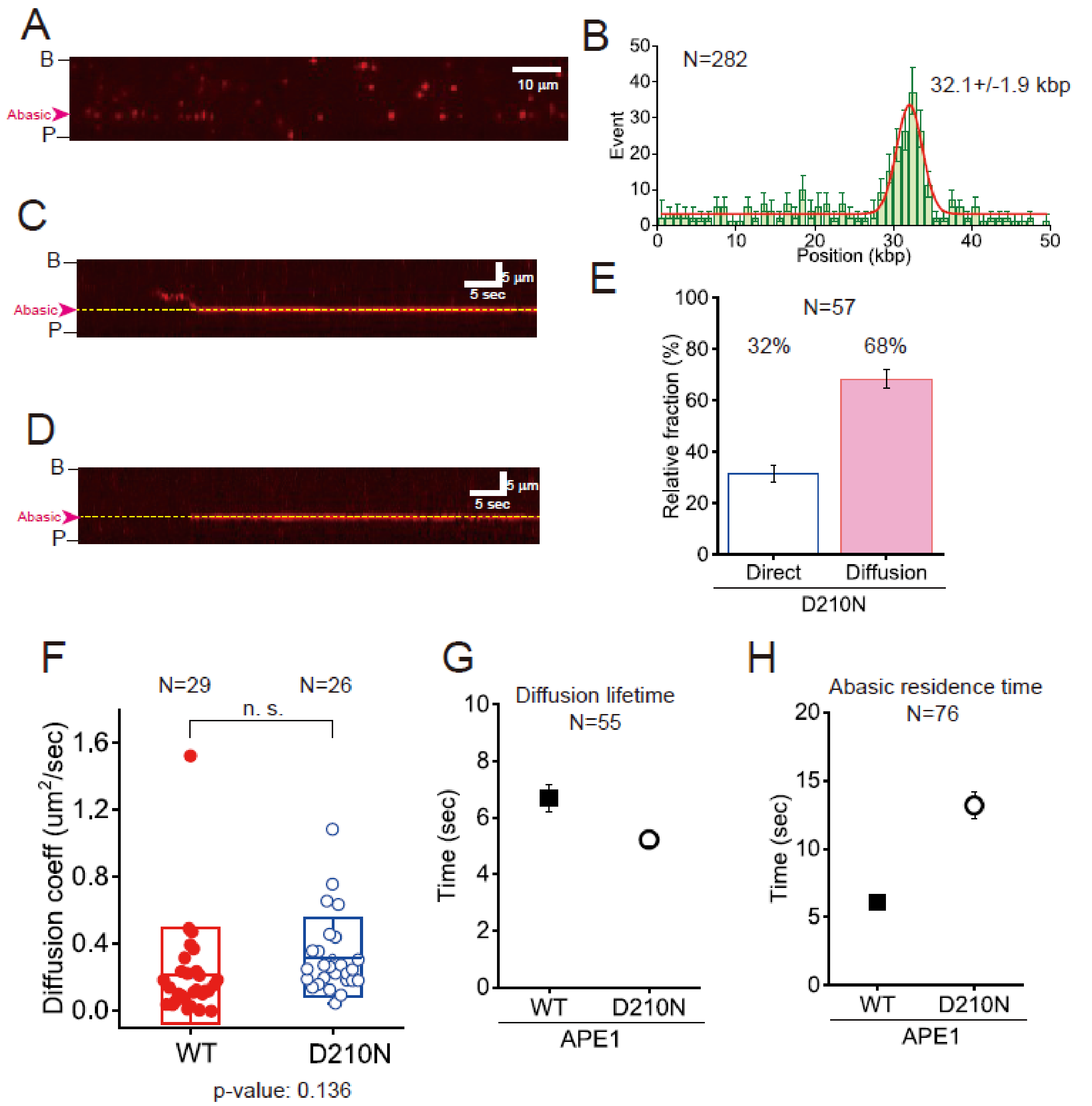


**Supplementary Figure 10. AP site search behavior of catalytic-dead APE1 mutant D210N.** (A) DNA curtain image of D210N binding to AP-containing λ-DNA. (B) Histogram of D210N binding positions, showing a peak at AP sites (~32.1 kb) with 70% confidence interval. (C, D) Kymographs of D210N searching by 1D diffusion (C) and 3D collision (D). (E) Relative fractions of 1D and 3D search modes for D210N. Error bars represent standard deviation from binomial distribution. n indicates total number of molecules analyzed. (F–H) Diffusion lifetime (F), AP site residence time (G), and D_diff (H) for wild-type and D210N APE1 at 100 mM NaCl and 0.1 mM $MgCl_2$. Student's t-test shows no significant difference in D_diff (P = 0.9977).

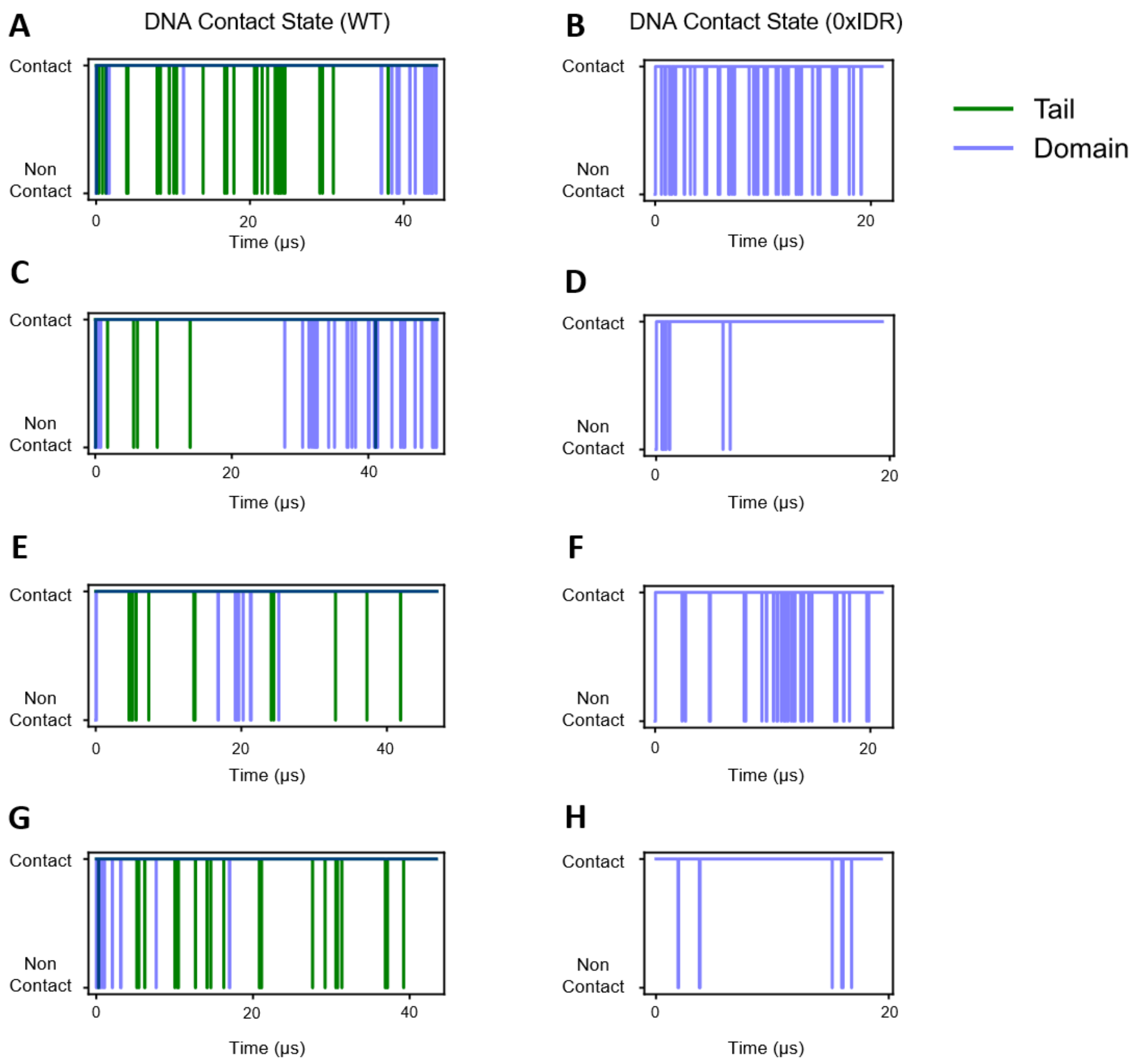


**Supplementary Figure 11. Time-resolved domain-level DNA-contact state for APE1.** A frame was labeled "in contact" if the distance from a residue's representative (terminal side-chain) atom to the DNA central axis was < 1.25 nm. The structured domain was considered "in contact" if at least one designated binding-site residue was in contact, and the IDR tail was considered "in contact" if any residue from positions 1–11 was in contact.

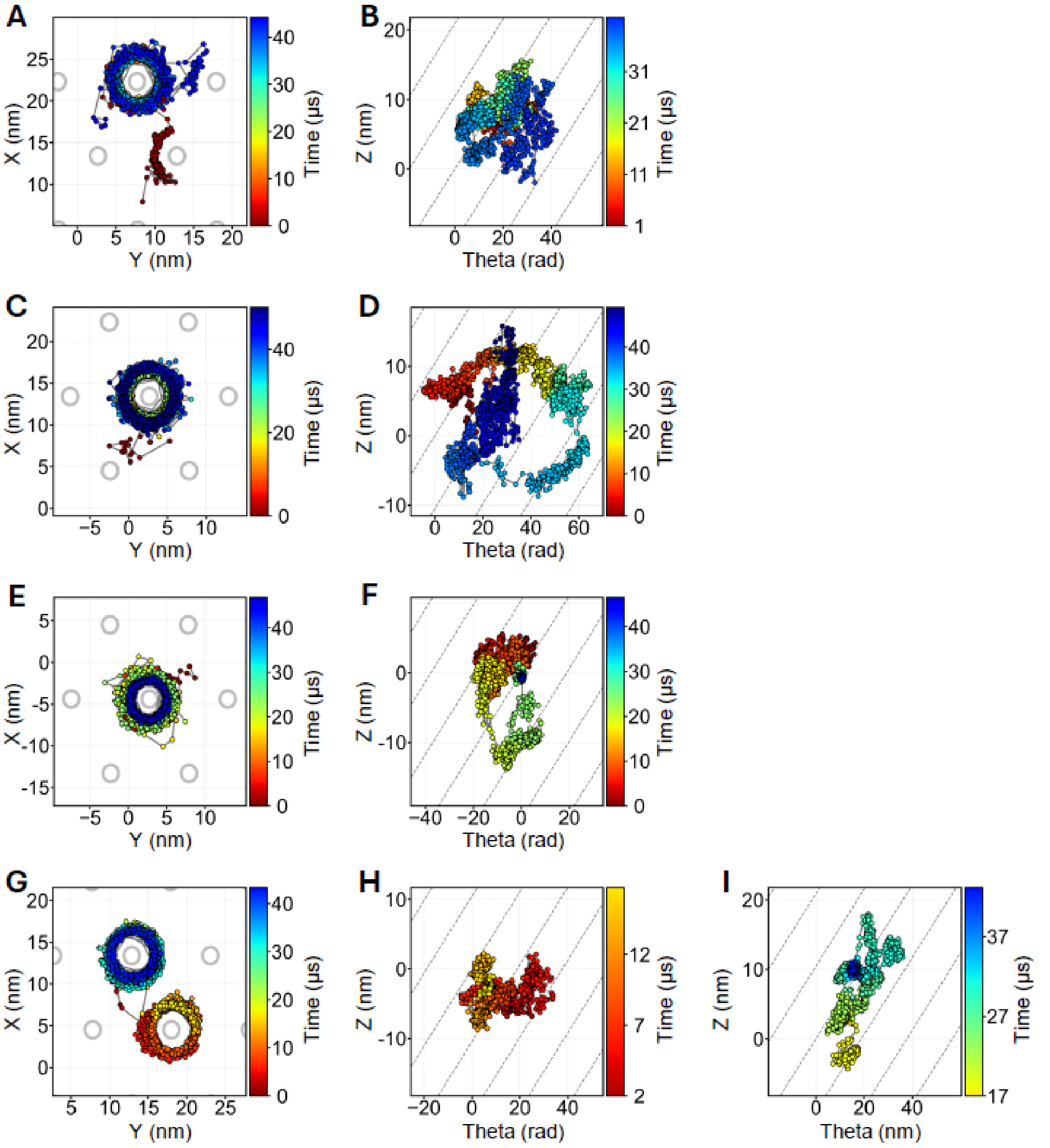


**Supplementary Figure 12. Center-of-mass (CoM) trajectories of wild-type APE1. (A, B)** CoM trajectories from simulation I projected onto the $xy$-plane (A) and the cylindrical $\theta - z$ space (B). Dashed lines in (B) represent the ideal helical trajectory expected for perfect groove-tracking motion, with a slope of $P/2\pi$, where $P = 3.4$ nm denotes the DNA helical pitch. Rainbow color scale indicates simulation time; gray circles mark the positions of periodic image DNA helices. **(C, D)** Same as panels A and B for simulation II. **(E, F)** Same as panels A and B for simulation III. **(G, H)** Same as panels A and B for simulation IV.

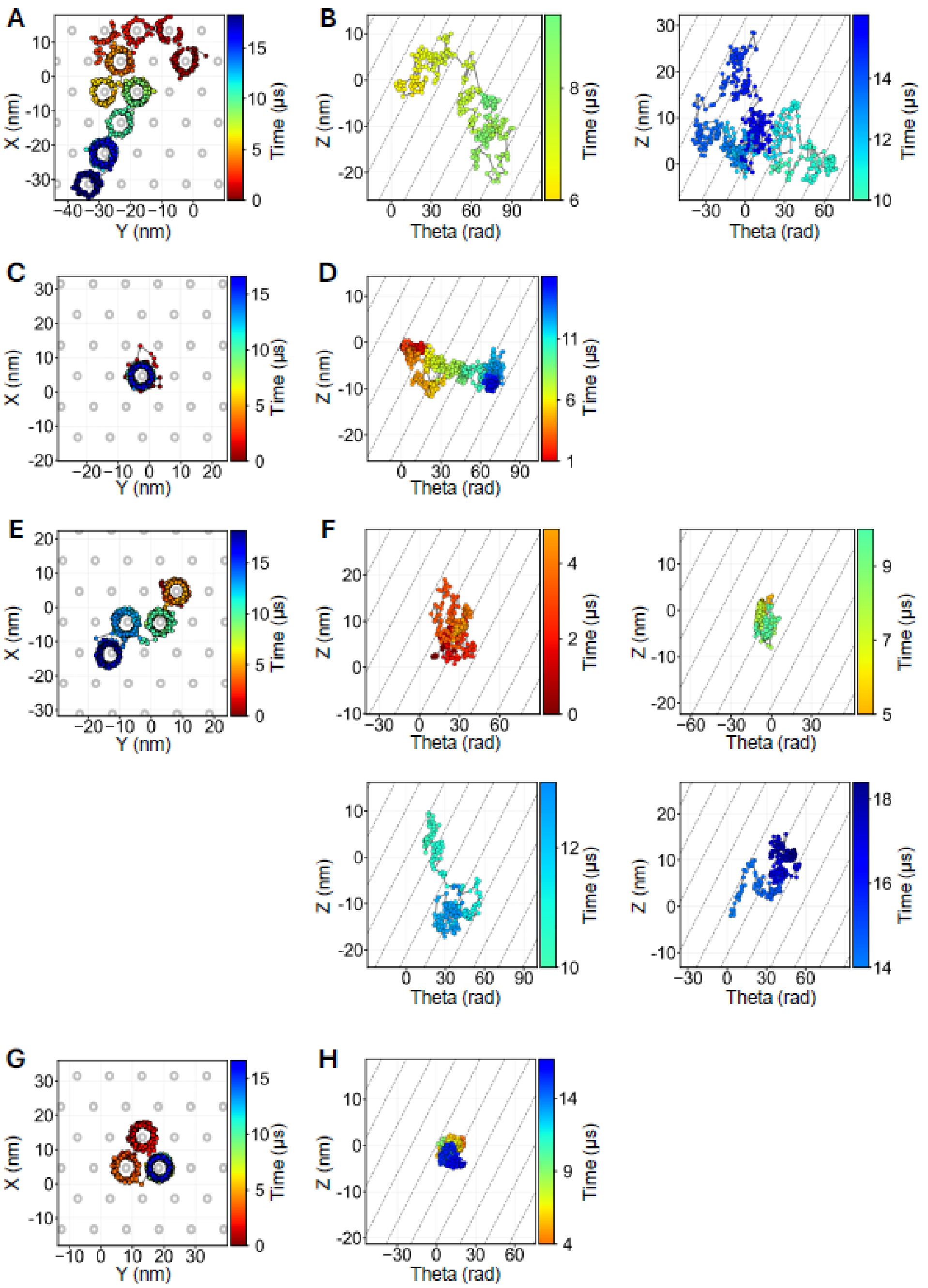


**Supplementary Figure 13. Center-of-mass (CoM) trajectories of the 0× IDR mutant. (A, B)** CoM trajectories from simulation I projected onto the $xy$-plane (A) and the cylindrical $\theta - z$ space (B). Dashed lines in (B) represent the ideal helical trajectory expected for perfect groove-tracking motion, with a slope of $P/2\pi$, where $P = 3.4$ nm denotes the DNA helical pitch. Rainbow color scale indicates simulation time; gray circles mark the positions of periodic image DNA helices. **(C, D)** Same as panels A and B for simulation II. **(E, F)** Same as panels A and B for simulation III. **(G, H)** Same as panels A and B for simulation IV.

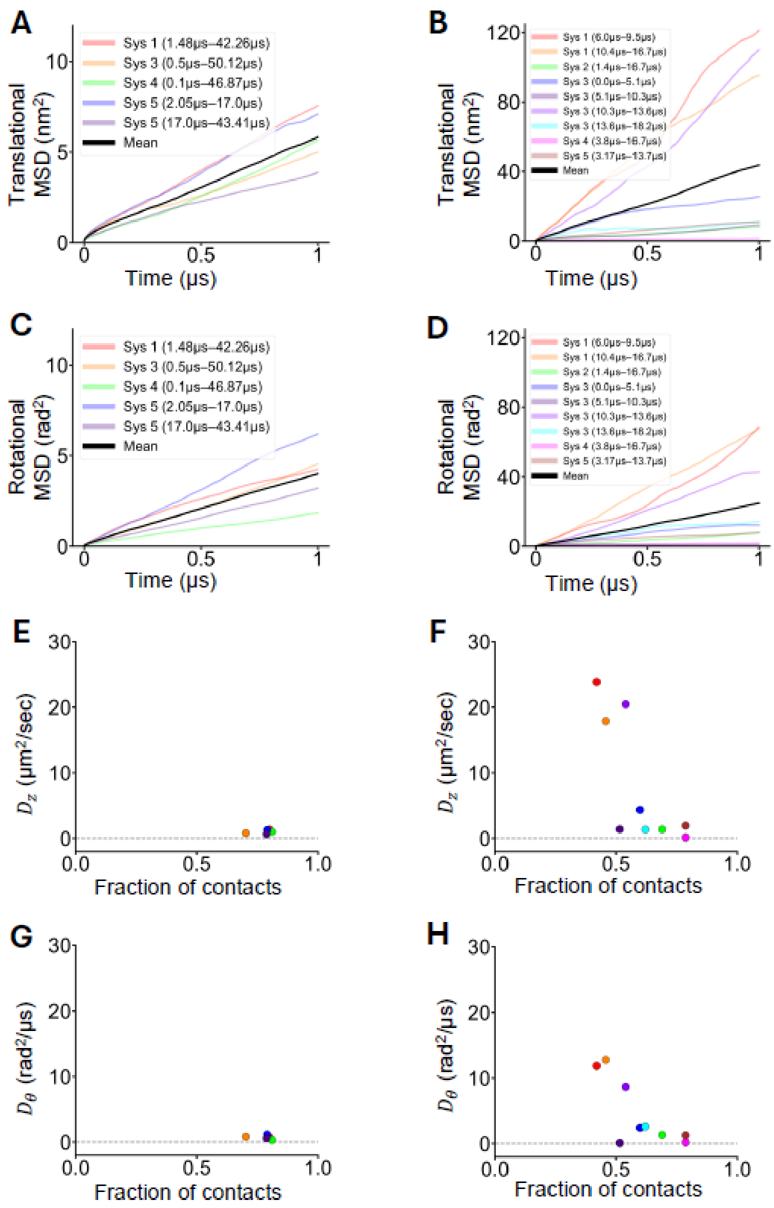


**Figure14. The correlation between the mean squared displacement (MSD) and the DNA contact fraction.** All simulations were segmented at each hopping event, so that every trajectory was divided into multiple intervals. For each interval, APE1's trajectory was realigned by centering on the nearest DNA, and only segments lasting at least 3 μs were retained to ensure the validity of the MSD analysis. (A, B) The translational MSD computed from the CoM axial positions for the wild-type APE1 (A) and the 0×IDR mutant (B), respectively. (C, D) The rotational MSD computed from the CoM angular orientations for the wild-type APE1 (C) and the 0×IDR mutant (D). (E) The X-axis denotes the DNA contact fraction averaged over all K/R residues in each segment of the wild-type APE1, and the Y-axis represents the translational diffusion coefficient along the DNA; both quantities were calculated for each of the previously defined intervals (as in A–D). (F) The corresponding translational diffusion coefficients for each segment of the 0×IDR mutant. (G, H) The rotational diffusion coefficient around the DNA axis versus the DNA contact fraction for each segment of the wild-type APE1 (G) and the 0×IDR mutant (H).

**Supplementary table I. Oligomer sequences.**

| Name | Sequence | Mfr. |
|---|---|---|
| Lambda R-biotin | 5'-[Phosphate]AGG TCG CCG CCC[biotin]-3' | Bioneer (South Korea) |
| Lambda L-biotin | 5'-[Phosphate]GGG CGG CGA CCT[biotin]-3' | Bioneer (South Korea) |
| Lambda L-Dig | 5'-[Phosphate]GGG CGG CGA[Dig] CCT[Dig]-3' | Bionics (South Korea) |
| Abasic lambda insertion oligo | 5'-[Phosphate] TGC ATG CGG CCG CTC TTC C[dSpacer]AT GGT GCG ATC GCT CTT CG -3' | Bioneer (South Korea) |
| Abasic DNA | 5'-[Cy5]GCGTCAAAATGT[spacer]GGTATTTCCATG-3' | Bionics (South Korea) |
| Abasic complementary DNA | 5'-CATGGAAATACCCACATTTTGACGC-3' | Bionics (South Korea) |
| 2AP-DNA AP strand | 5'- GGT GGT GGT AAG ATG ATG AAG AGA AC[dSpacer] GTG CGT CGG AGA GAA TTC GGA GAG AGA AC[dSpacer] GTG CGT GTG GCG-3' | Integrated DNA Technologies (USA) |
| 2AP-DNA biotin strand | 5'- CGC CAC ACG CAC TGT TCT CTC TCC GAA TTC TCT CCG ACG CAC TGT TC[iAmC6T] CTT CAT CAT CTT ACC ACC ACC[biotin] -3' | Integrated DNA Technologies (USA) |
| Undamaged DNA strand | 5'-GGT GGT GGT AAG ATG ATG AAG AGA ACA GTG CGT CGG AGA GGA GAG AGA GAA GGA AGT GTG GCG GAA GG-3' | Integrated DNA Technologies (USA) |
| AP-DNA AP strand | 5'- GGT GGT GGT AAG ATG ATG AAG AGA ACA GTG CGT C[dSpacer]G AGA GGA GAG AGA GAA GGA AGT GTG GCG GAA GG-3' | Integrated DNA Technologies (USA) |
| Undamaged/AP-DNA complementary strand | 5'- CGC CAC ACT TCC TTC TCT CTC TCC TCT CCG ACG CAC TGT TC[iAmC6T] CTT CAT CAT CTT ACC ACC ACC[biotin] -3' | Integrated DNA Technologies (USA) |
